# The Pick fold in tau filaments from human *MAPT* mutants

**DOI:** 10.64898/2026.03.08.710379

**Authors:** Chao Qi, Sofia Lövestam, Jenny Shi, Alexey G. Murzin, Sew Peak-Chew, Thomas T. Warner, Harro Seelaar, Patrick W. Cullinane, Zane Jaunmuktane, John C. van Swieten, Sjors H.W. Scheres, Michel Goedert

## Abstract

Mutations in *MAPT*, the tau gene, give rise to forms of frontotemporal dementia and parkinsonism linked to chromosome 17 (FTDP-17T), with abundant filamentous tau inclusions in brain cells. Some mutations that encode missense and deletion variants can give rise to a clinical picture of Pick’s disease and filaments made of three-repeat tau. Here we report the electron cryo-microscopy (cryo-EM) structures of tau filaments from individuals with *MAPT* mutations D252V, G272V, S320F and ΔG389-I392. The two-layered Pick fold was present in the individuals with mutations D252V and ΔG389-I392. By contrast, mutations G272V and S320F gave rise to a more open variant of the Pick fold, with residues 272-341 rotated by 20-25° with respect to the rest of the structure. These findings show that missense mutations within the filament core can modify the Pick fold, generating closely related structural variants. In addition, we were able to reconstitute the Pick fold and some of its variants using seeded assembly with recombinant 0N3R tau carrying 12 serine or threonine to aspartate substitutions (PAD12) and missense mutations D252V, G272V or S320F. This work provides a foundation for the development of structure-based diagnostic and therapeutic approaches.

## Introduction

In the adult human brain, six tau isoforms are expressed from a single *MAPT* gene by alternative mRNA splicing (1). They differ by the inclusion or exclusion of two exons near the N-terminus (exons 2 and 3) and a single exon near the C-terminus (exon 10) of the protein. Exon 10 encodes a repeat of 31 amino acids, and its inclusion gives rise to three isoforms with four repeats (4R). The other three isoforms lack exon 10 expression and have three repeats (3R). These repeats and adjoining sequences constitute the microtubule-binding domains of tau. Part of this sequence also forms the core of assembled tau in neurodegenerative diseases, indicating that the physiological function of microtubule binding and the pathological assembly into amyloid filaments are mutually exclusive (2).

Mutations in *MAPT* cause inherited forms of frontotemporal dementia and parkinsonism linked to chromosome 17 (FTDP-17T) (3-5) with abundant filamentous tau inclusions in brain cells that are made of either 3R, 4R or 3R+4R tau. These mutations influence the alternative mRNA splicing of exon 10 and/or the ability of tau protein to interact with microtubules, resulting in tau filament formation (6). Mutations that cause an overproduction of wildtype 3R or 4R tau result in the deposition of 3R tau with the Pick fold (7) or 4R tau with the argyrophilic grain disease (AGD) fold (8). Filamentous inclusions of 3R+4R tau associated with missense mutations V337M and R406W adopt the Alzheimer fold (9). By contrast, 4R tauopathies caused by *MAPT* missense mutations P301L and P301T give rise to distinct folds (10). Missense mutations P301L, P301T, V337M and R406W reduce the ability of tau to interact with microtubules. Filaments from a case with missense mutation S305I, that has effects at both mRNA and protein levels, also adopt a unique tau fold (11).

A key priority is to develop model systems that replicate the same tau filament structures as those observed in disease, which may allow one to identify the causes underlying the structural specificity of amyloid filaments (2). This has previously only been achieved for the Alzheimer and the chronic traumatic encephalopathy folds of assembled tau (12-14).

Here we present the cryo-EM structures of tau filaments extracted from the frontal and temporal cortices of FTDP-17T individuals with tau mutations D252V, G272V and ΔG389-I392, as well as from the caudate nucleus of an individual with tau mutation S320F. The Pick fold, which has previously been identified in tau filaments extracted from the frontotemporal cortex of an individual with sporadic Pick’s disease (15) and two individuals with *MAPT* mutation DK281 (7) was present in D252V and ΔG389-I392 filaments, whereas a more open Pick fold was present in G272V and S320F filaments. When using tau seeds from these cases of FTDP-17T and recombinant 0N3R PAD12 tau with mutations D252V, G272V or S320F, we were able to form the Pick fold and some of its variants.

## Results

### Structures of tau filaments from individuals with the *MAPT* mutations encoding D252V or ΔG389-I392 tau

Individuals with familial frontotemporal dementias carrying the heterozygous *MAPT* mutations encoding D252V or ΔG389-I392 were reported in 2019 (16). We determined the cryo-EM structures of tau filaments extracted from the frontal cortex of these two cases to resolutions of 2.8 Å and 2.3 Å, respectively (Figures 1 and 2).

**Figure 1.**
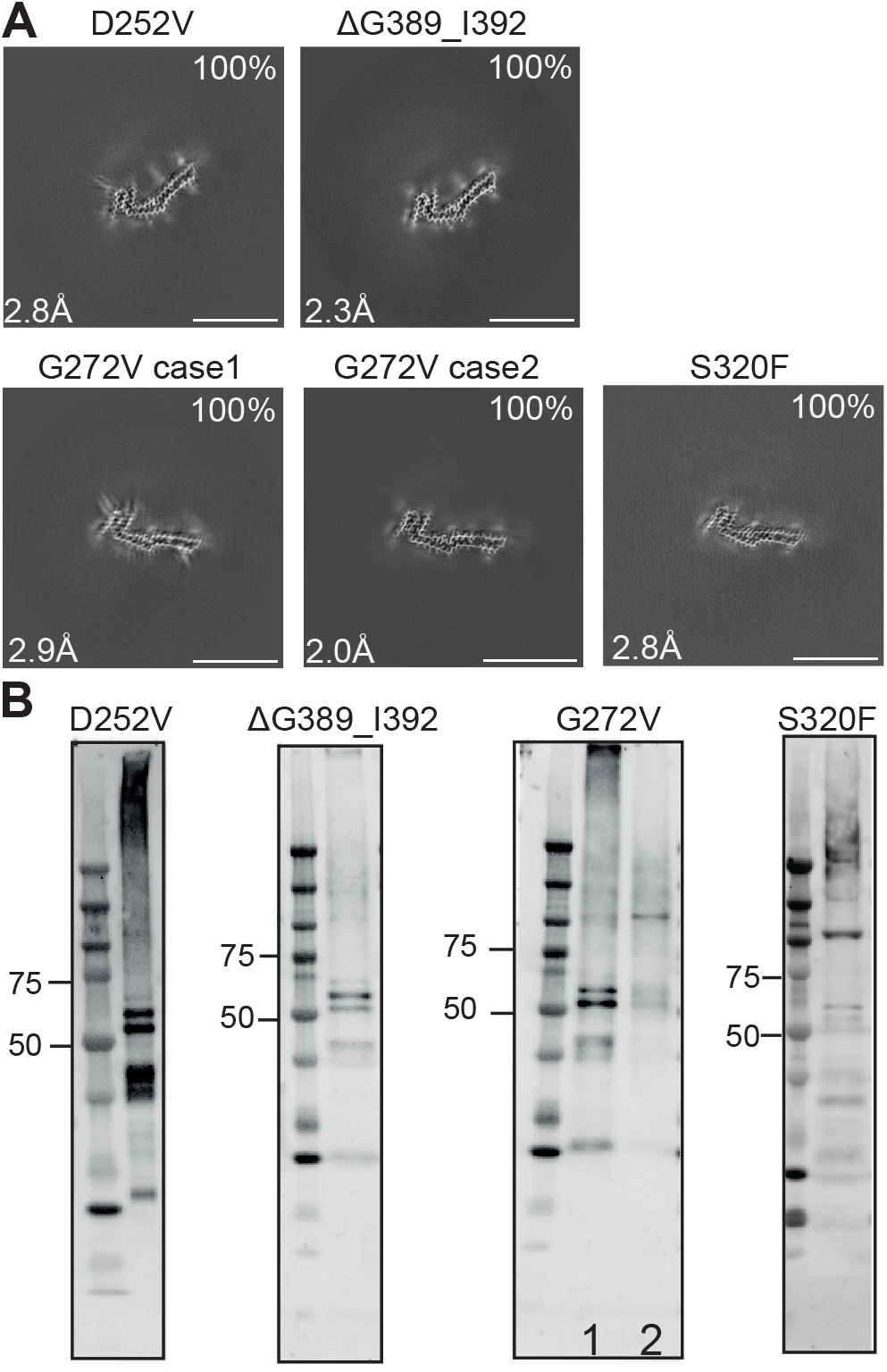
Mutations encoding D252V, ΔG389_I392, G272V and S320F in *MAPT*: cryo-EM cross-sections of tau filaments and immunoblot analysis. **A**, Cryo-EM cross-sections of tau filaments perpendicular to the helical axis, each representing approximately one rung in projected thickness. The structural resolutions (in Å) and percentages of filament types (%) are indicated at the bottom left and top right, respectively. Scale bar, 10 nm. **B**, Immunoblotting of sarkosyl-insoluble tau from individuals with MAPT mutations encoding D252V, ΔG389_I392, G272V and S320F. Phosphorylation-independent anti-tau antibody BR134 was used for D252V and ΔG389_I392, G272V; anti-tau antibody BR133 was used for S320F.

**Figure 2.**
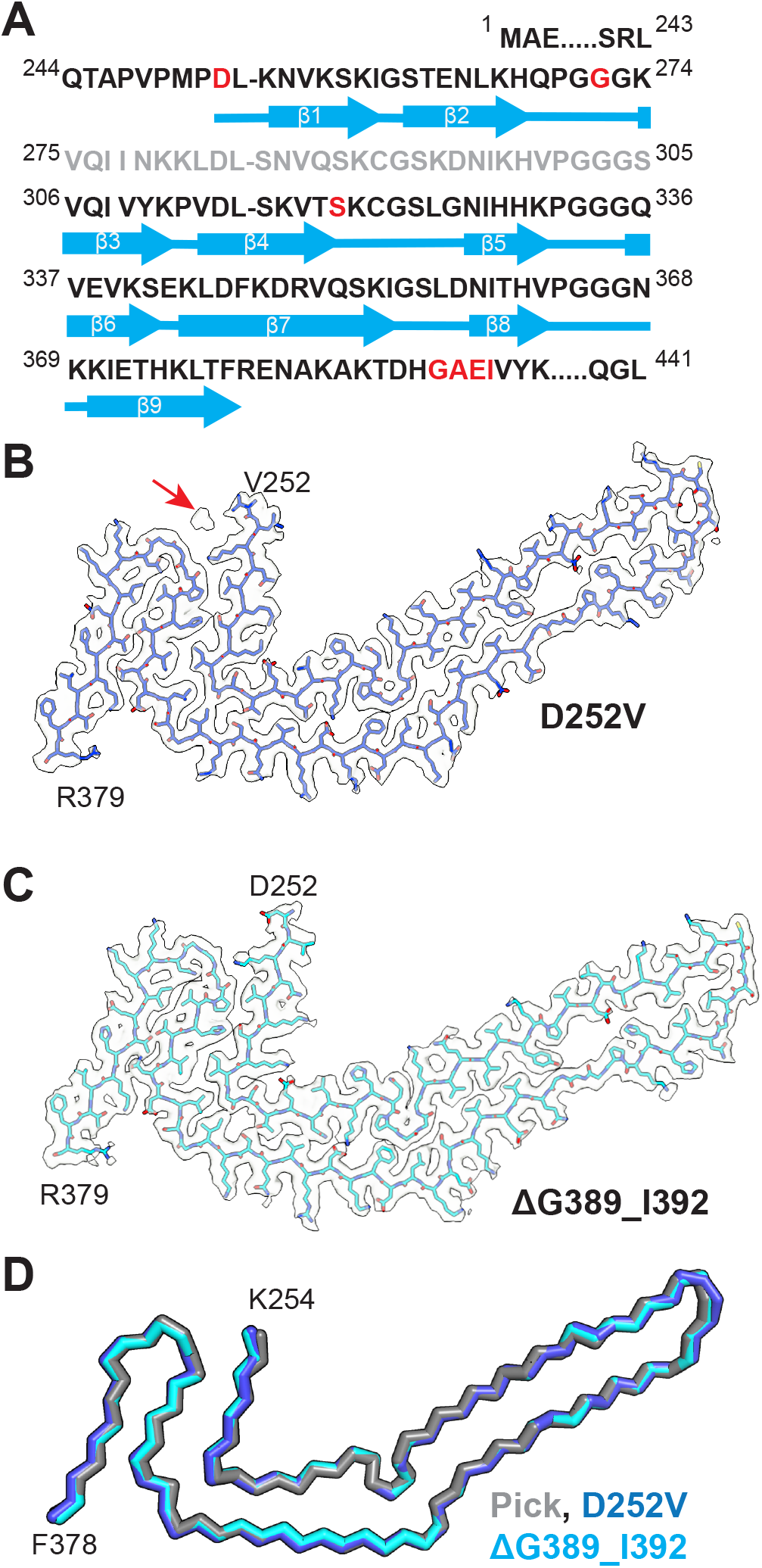
Cryo-EM structures of tau filaments from the frontal cortex of individuals with *MAPT* mutation D252V and ΔG389_I392. **A**, Amino acid sequence of repeats R1–R4 of tau (residues 244–368) and the sequence after R4. The core of the tau filaments extends from residues D252/V252 to R379, with the exclusion of R2. It consists of nine β-strands (β1–β9, thick arrows) connected by loops (thin lines). The positions of the D252V, G272V, S320F and ΔG389_I392 mutations are highlighted in red. **B**, Cryo-EM density map and atomic model of the tau filament with the D252V mutation, which adopted the Pick fold. The red arrow indicates the cofactor near V252. **C**, Cryo-EM density map and atomic model of the tau filament with the ΔG389_I392 mutation, which also adopted the Pick fold. Deleted residues G389_I392 are located out-side the ordered core. **D**, Overlay of tau filament structures with D252V (blue), ΔG389_I392 (cyan) and the Pick fold (grey, PDB 6GX5). The RMSD between Cα atoms of D252V and the Pick fold was 0.695 Å, and that between Cα atoms of ΔG389_I392 and the Pick fold was 0.389 Å.

The D252V mutation in tau was identified in a male who developed symptoms of frontotemporal dementia (FTD) at age 46 and died aged 54. He presented with behavioural symptoms and language difficulties and showed signs of memory impairment. There was a family history of dementia, with the proband’s mother developing dementia in her 70s and all but one of her seven siblings also developing dementia. Neuropathologically, there was global atrophy that was most severe in the frontal and temporal lobes. Tau immunohistochemistry revealed the presence of abundant 3R tau-immunoreactive Pick bodies, especially in cortical grey matter. Western blotting of sarkosyl-insoluble tau from frontal cortex showed the presence of two strong bands of 60 and 64 kDa and a weak band of 68 kDa (Figure 1B).

The ΔG389-I392 deletion in tau was identified in a 49 year-old female with a two-year history of behavioural changes and decline in language function who died aged 54. The patient’s mother died in middle age without a diagnosis of dementia, but with a history of aggressive behaviour. Her mother’s mother developed dementia with behavioural changes in her 50s and died aged 70. Neuropathologically, there was severe frontal and medial temporal lobe atrophy, with abundant 3R tau-immunoreactive Pick bodies. Western blotting of sarkosyl-insoluble tau from frontal cortex showed the presence of two strong bands of 60 and 64 kDa and a weak band of 68 kDa (Figure 1B).

Frontal cortex from the D252 and ΔG389-I392 cases only contained tau filaments made of a single protofilament that were identical in structure to each other and to the previously determined Pick fold (7,15) (Figure 2A-C). Superpositions onto the Pick structure (PDB:6GX5) yielded root-mean-square deviation (RMSD) values of 0.695 Å (D252V) and 0.389 Å (ΔG389-I392) between Cα atoms of the ordered core residues K254-F378, indicating structural identity. The quality of our maps allowed modelling of two additional residues (D/V252 and L253) at the amino-terminus and one additional residue (R379) at the carboxy-terminus of the Pick tau fold. Similar to wild-type tau filaments, there were external non-proteinaceous densities in tau filaments from both mutants; they were present near positively charged and non-polar side chains.

In the D252V tau map, the density at the side chain position of residue 252 was bulkier than in the ΔG389-I392 map and a disconnected density was present between this residue and the 364PGGG367 motif in R4, indicative of the presence of the mutant V252 residue in the filaments (Figures 2B, S2A). Residues G389-I392 lie outside the ordered filament core. Therefore, the cryo-EM maps cannot provide any insight into whether the tau filaments are made of wild-type or mutant tau. Mass spectrometric analysis identified peptides corresponding to both wild-type and mutant tau in the D252V and ΔG389-I392 frontal cortical extracts (Figure S1), indicating co-assembly of wild-type and mutant human tau into filaments with the Pick fold.

### Structures of tau filaments from individuals with the *MAPT* mutation encoding G272V tau

Dominantly inherited Pick’s disease is caused by the G272V mutation in *MAPT* (4,17-22). Fresh-frozen temporal cortex from two individuals carrying this mutation has been characterised (21,22). We determined the cryo-EM structures of tau filaments extracted from the temporal cortex of both individuals to resolutions of 2.9 Å and 2.0 Å (Figures 1 and 3).

**Figure 3.**
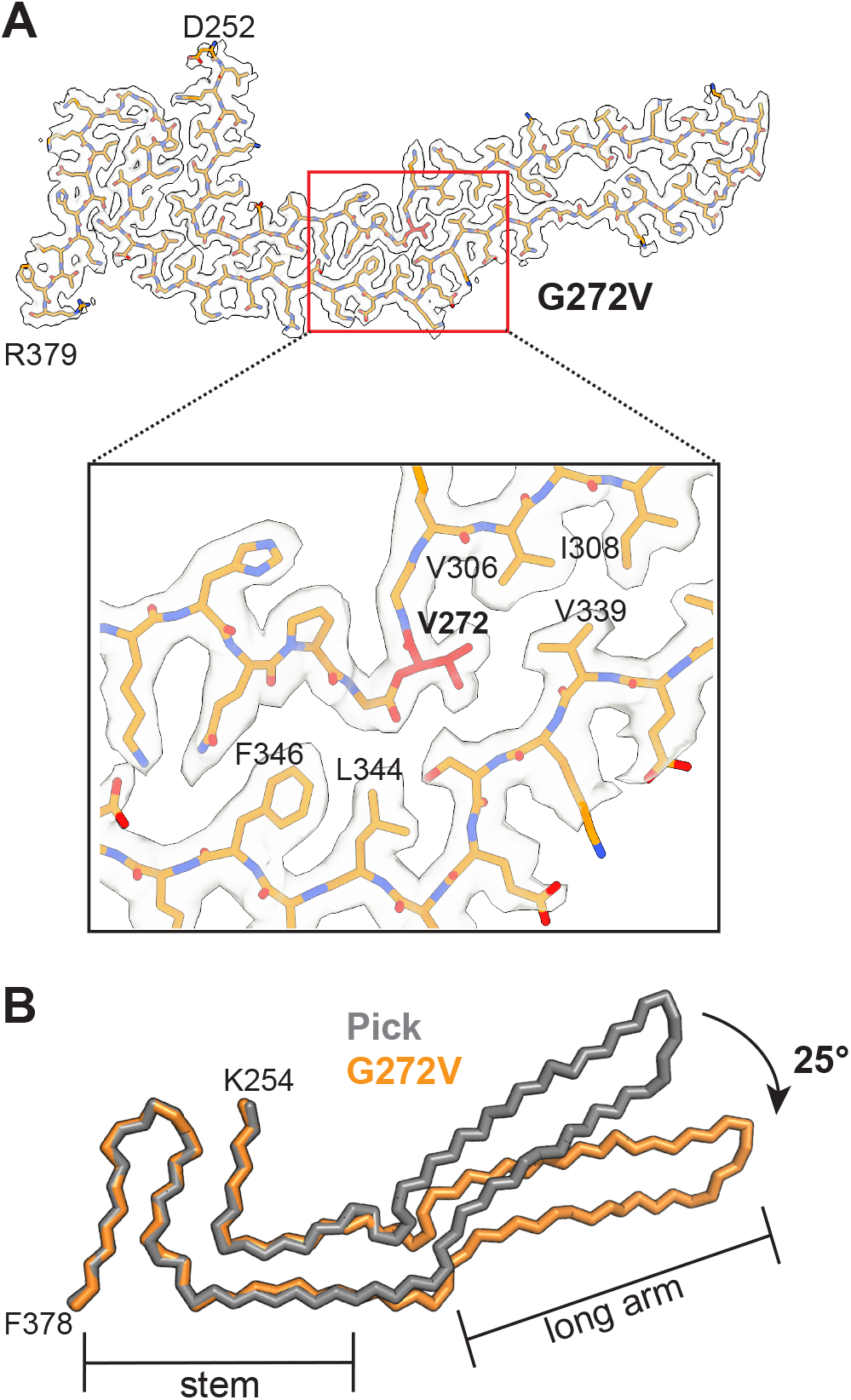
Cryo-EM structure of tau filaments from the temporal cortex of case 2 with *MAPT* mutation G272V. **A**, Cryo-EM density map and atomic model of tau filaments with the G272V mutation. Residue V272 is shown in red. *Inlet*: enlarged view of the V272 region, where the side chain of V272 inserts into a hydrophobic pocket formed by residues V306 and V339. **B**, Overlays of tau filament structures with G272V (orange) and the Pick fold (grey, PDB 6GX5), aligned using the stem region. The long arm region in the G272V filament structure is rotated by approximately 25° relative to the Pick fold.

Case 1 developed symptoms of FTD at age 45 and died aged 54; case 2 developed symptoms of FTD at age 52 and died aged 67. Both brains showed severe neuronal loss in many regions, including the temporal cortex, accompanied by abundant 3R tau-positive Pick bodies that were not phosphorylated at S262. Pick bodies in sporadic Pick’s disease are also not phosphorylated at S262 (23). By Western blotting, we identified strong abnormal tau bands of 60 and 64 kDa and a weak band of 68 kDa (Figure 1B), in confirmation of previous findings (21).

Tau filaments were made of a single protofilament, with identical ordered cores that extended from residues D252-R379. They revealed a more open variant of the Pick fold with essentially the same secondary structures (Figure S3). Separate superpositions of the stem (β1-2, β7-β9) and the long arm regions (β3-β6) of the Pick fold (15; PDB:6GX5) yielded RMSD values of 0.408 Å and 0.476 Å, respectively (Figure S3). Residue V272 lies within the ^269^QPGGG^273^ motif at the end of R1 (Figure 2A), where its hydrophobic side chain inserts into a pocket formed by V306, I308 and V339 of tau (Figure 3A), thus altering the local conformation of the motif and propagating a global rearrangement. Compared to the canonical Pick fold (15), the side chains of tau residues Q269 and P270 switched their in-out orientations in the G272V variant, whereas the long arm rotated by about 25° relative to the stem (Figure 3B).

Although the G272V structure determined at 2.0 Å resolution had a clearly resolved V272 side chain, its density itself could not rule out co-assembly of wild-type (G272) and mutant (V272) tau. Mass spectrometry did show the presence of peptides containing both variants (Figure S4). An additional surface density was present near H268 and K274 of the G272V tau filament that was not observed in the wild-type Pick fold (Figure S2). The identity of this density, which may represent a post-translational modification of tau or negatively charged molecules that countered the positive charge of K274, remains unknown.

### Structures of tau filaments from an individual with the *MAPT* mutation encoding S320F tau

The dominantly inherited S320F mutation in tau was identified in a male who developed symptoms of FTD at the age of 38 and died aged 53 (24). We determined the cryo-EM structures of tau filaments extracted from the caudate nucleus of the S320F case to a resolution of 2.8 Å (Figures 1 and 4).

**Figure 4.**
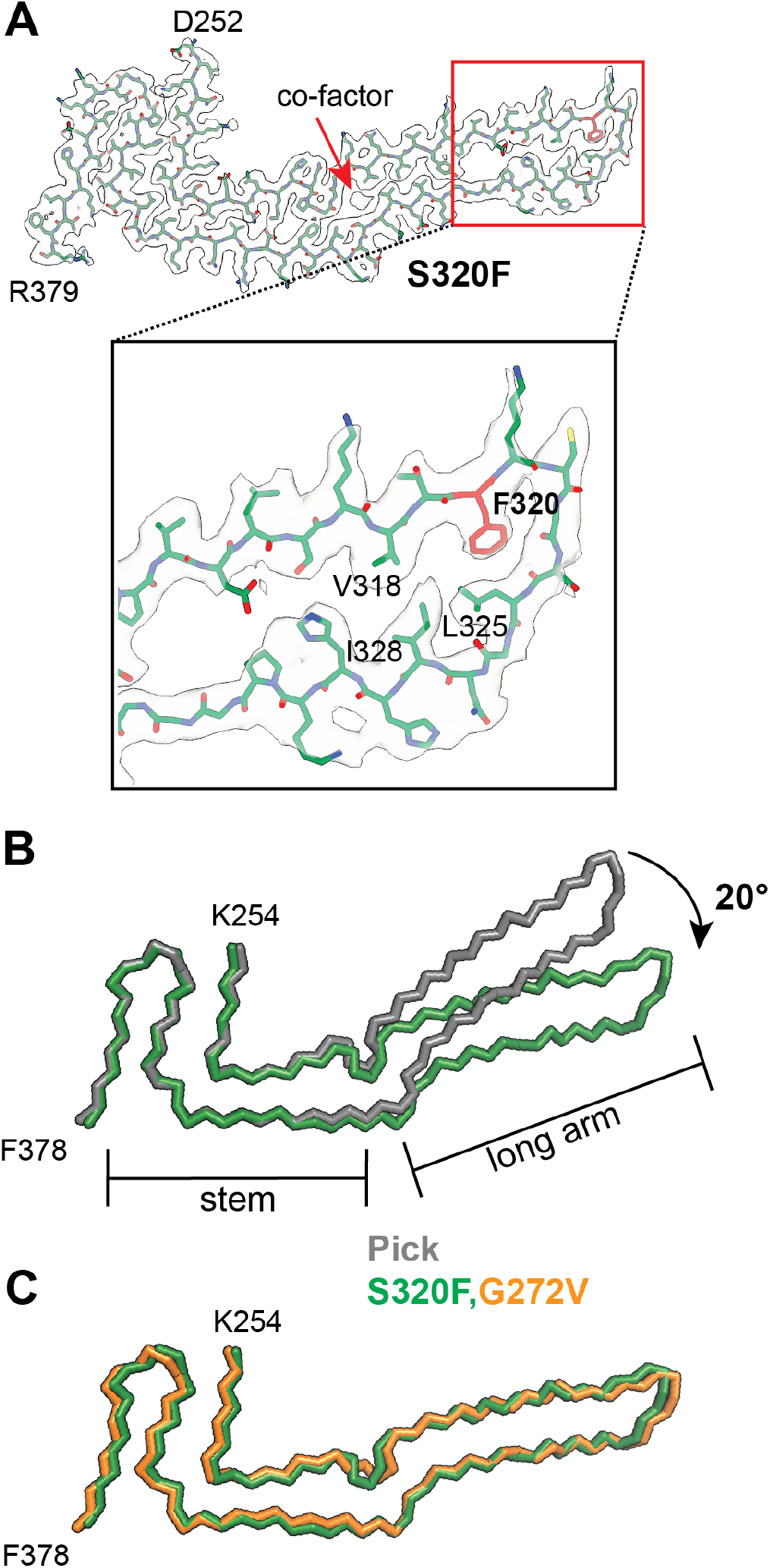
Cryo-EM structure of tau filaments from the caudate nucleus of the individual with *MAPT* mutation S320F. **A**, Cryo-EM density map and atomic model of tau filaments with the S320F mutation. Residue F320 is shown in red. *Inlet*: enlarged view of the F320 region showing that the phenylalanine side chain points into a hydrophobic pocket formed by residues V318, L325 and I328. **B**, Over-lay of tau filament structures with S320F (green) and the Pick fold (grey, PDB 6GX5), aligned using the stem region. The long arm region in the S320F filament structure is rotated by approximately 20° relative to the Pick fold. **C**, Structural overlay of the S320F (green) and G272V (orange). The RMSD between Cα atoms of both structures was 1.4 Å.

The proband’s mother died with a similar dementing illness. Neuropathologically, there was severe frontal and medial temporal lobe atrophy, with abundant 3R tau-immunoreactive Pick bodies. By Western blotting of sarkosyl-insoluble tau from the frontal cortex, two strong bands of 60 and 64 kDa and a weak band of 68 kDa were present (Figure 1E), in confirmation of previous findings (24).

All filaments consisted of a single protofilament. Like tau filaments extracted from the temporal cortex of individuals with the *MAPT* mutation encoding G272V tau, the S320F tau filaments adopted a more open conformation relative to the Pick fold. Separate superpositions of the stem and long-arm regions yielded RMSD values of 0.603 Å and 0.599 Å (Figure S3B), with a relative rotation of the long arm region by about 20° relative to the Pick fold (Figure 4B). The side chain density of residue 320 was significantly larger than that of a serine, indicating the presence of mutant F320 in the tau filaments. The side chain of F320 inserts into a hydrophobic pocket formed by residues V318, L325 and I328, which is made possible by a slight adjustment of the side chain conformations of those residues. Mass spectrometric analysis of the sarkosyl-insoluble fractions from the temporal cortex detected peptides containing S320 and F320 (Figure S4C), indicating the co-assembly of wild-type and mutant tau in the S320F filaments.

It is not clear why the S320F tau filament core is more open than the canonical Pick fold and why it resembles the G272 variant fold. There are local conformational changes in the 269QPGGG273 motif, but without switching the in-out orientations of the side chains of residues Q269 and P270. A small non-proteinaceous density was present between G272 and the hydrophobic side chains of V306 and V339 (Figure 4A). Its position overlapped with that of the side chain of V272 in the G272V tau filament structure. Although the molecular identity of this density is not known, it may mimic the side chain of valine and stabilise a G272V-like conformation. Like the G272V tau filaments, the S320F tau filaments had an additional density near H268 and K274 (Figure S2D).

### Seeded aggregation *in vitro* to assemble the Pick tau fold and variant filaments

We initially performed seeded aggregation *in vitro* with recombinant 0N3R PAD12 tau, using seeds extracted from the frontal cortex of the individual with sporadic Pick’s disease that was used for the identification of the Pick tau fold (15). A minority of filaments had two protofilaments (doublets) with the Pick fold (Figure 5A), but most filaments were prone to untwisting, precluding cryo-EM structure determination.

**Figure 5.**
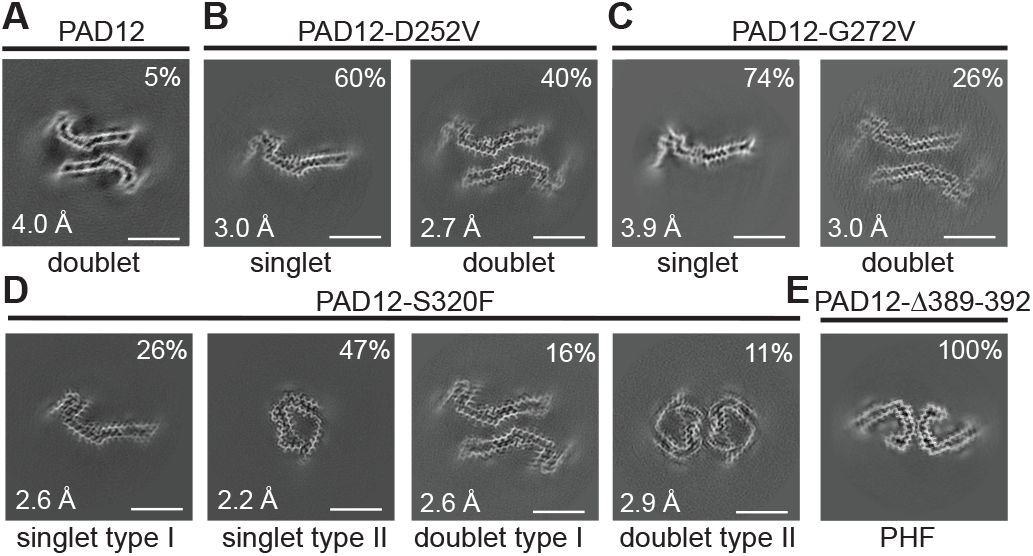
Seeded assembly experiments with mutations D252V, G272V, S320F and ΔG389-I392 in PAD12 tau. Cryo-EM cross-sections of tau filaments assembled from: **A**, PAD12 and seeds from the case of sporadic Pick’s disease. **B**, PAD12-D252V and seeds from the case with *MAPT* mutation D252V. **C**, PAD12-G272V and seeds from case 2 with *MAPT* mutation G272V. **D**, PAD12-S320F and seeds from the case with *MAPT* mutation S320F. **E**, PAD12-ΔG389-I392 and seeds from the case with *MAPT* mutation ΔG389-I392. The structural resolutions (in Å) and percentages of filament types (%) are indicated at the bottom left and top right, respectively. Scale bar, 5 nm. No tau seeds were detected during cryo-EM processing.

Given that the tau filaments from the FTDP-17T cases described above adopted the Pick fold or variants thereof, we also performed seeded aggregation *in vitro* using recombinant PAD12 tau with an additional D252V, G272V, S320F or ΔG389-I392 mutation. The assembly of recombinant PAD12 tau with one of these mutations into filaments was induced by the addition of 1 µl of brain-derived filaments (corresponding to 10 µg of brain tissue) to 30 µl of recombinant tau.

We then determined the cryo-EM structures of the seeded tau aggregates (Figure 5).

Seeding PAD12-D252V recombinant tau with tau seeds extracted from the frontal cortex of the individual with the D252V mutation yielded singlets (one protofilament) and doublets (two protofilaments) (Figures 5 and 6). The ordered cores of the protofilaments spanned residues 255-363, adopting the same conformation as in the Pick fold (Figure 7). Residues 364-379, which form a hairpin conformation in the Pick fold, were less ordered in the recombinant structure, with some residual densities being present in the map (Figure 5B). The doublets were stabilised by electrostatic interactions at the protofilament interface, with K340 and K343 from one protofilament forming salt bridges with E342 and E338 from the opposing protofilament and vice versa (Figure 6A).

**Figure 6.**
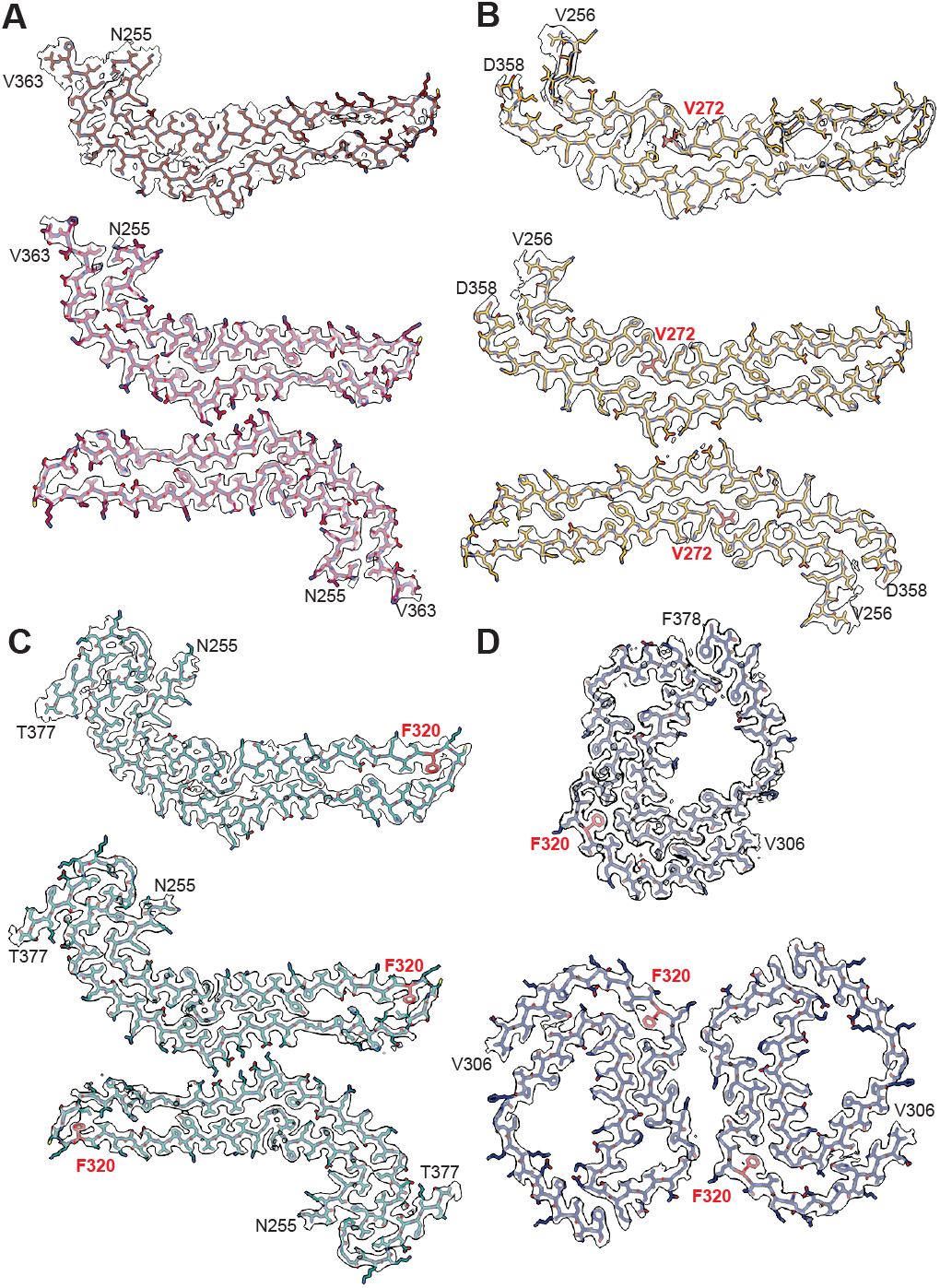
Cryo-EM density maps and atomic models of recombinant PAD12 tau filaments with mutations D252V, G272V and S320F. **A**, D252V. **B**, G272V. **C**, S320F Pick fold. **D**, S320F type 2.

**Figure 7.**
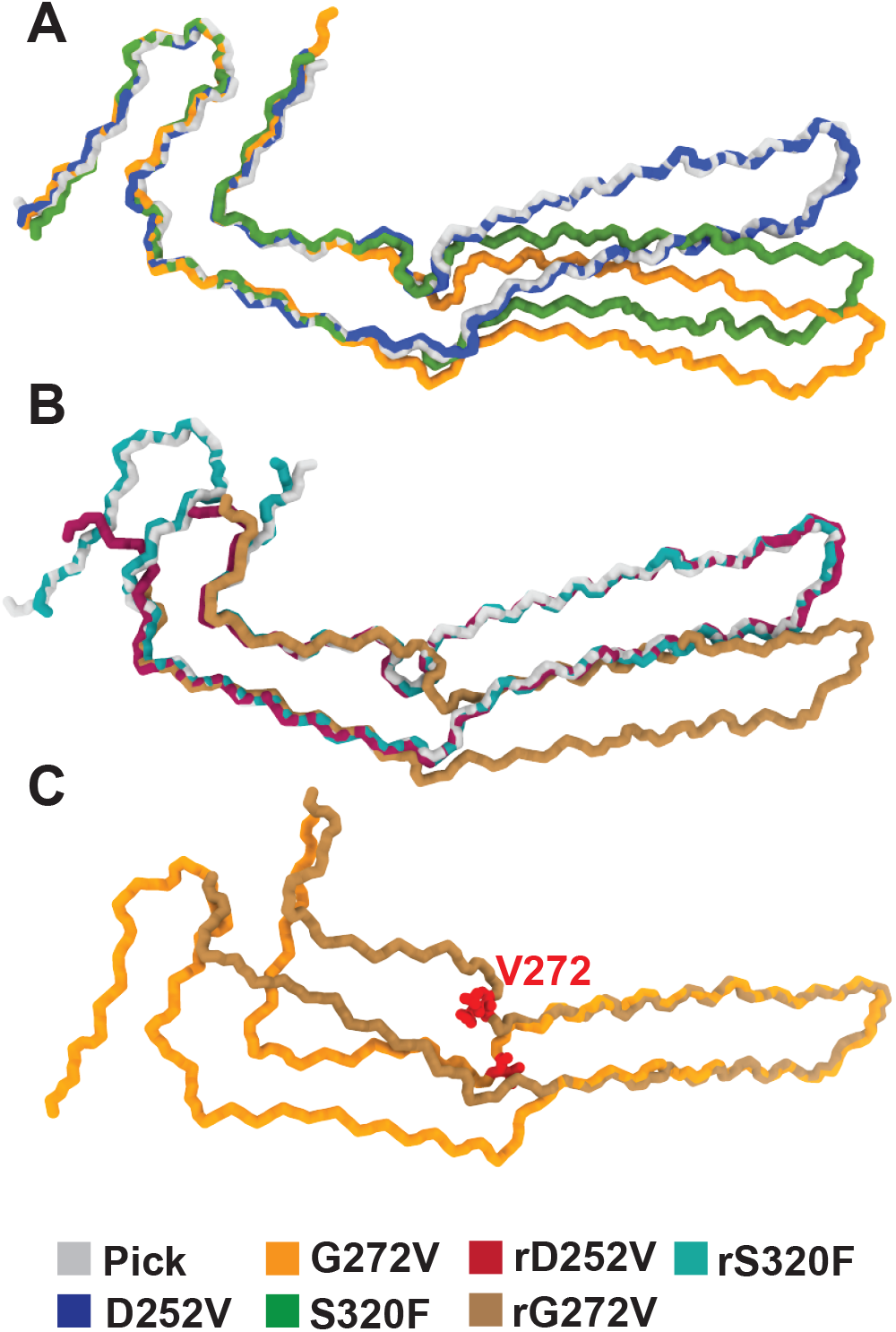
Comparison of the structures of tau filaments from individuals with FTDP-17T mutations (D252V, G272V and S320F) with those of filaments from an individual with sporadic Pick’s disease. **A,** Structural overlay of the Pick fold (grey) with D252V (blue), G272V (orange) and S320F (green). B, Structural overlay of the Pick fold (grey) and recombinant (r) PAD12 tau filaments with mutations D252V (maroon), G272 (brown) and S320F (teal). C, Comparison of the structures of G272V *ex vivo* filaments with those formed using seeded assembly (V272 is shown in red).

Seeding PAD12-G272V recombinant tau with seeds extracted from the temporal cortex of case 2 with the G272V mutation (21) yielded both singlet and doublet filament subtypes, the ordered cores of which spanned residues (256-358); they also showed the same interactions across the entire structures, except for the V272 mutation site. The 269QPGGG273 motif adopted a different conformation compared to that of the G272V filaments from the human brain, with V272 forming new interactions with L344 and F346. Like in G272V brain tau filaments, the side chains of residues Q269 and P270 switched their in-out orientations compared to the canonical Pick fold (15). This resulted in different relative orientations of the stem and long arm substructures compared to those in brain-derived Pick folds of wildtype tau and the G272V mutant (Figure 7).

Seeding PAD12-S320F recombinant tau with seeds extracted from the caudate nucleus of the individual with the S320F mutation yielded filaments with four different structures. One type of singlet and one type of doublet consisted of protofilaments with the canonical Pick fold, including the hairpin conformation at residues 359-377. Interestingly, in these filaments the phenylalanine side chain at residue 320 did not induce the open variant of the Pick fold that was observed in brain-derived filaments. The other two filament types were a singlet and a doublet with a different fold and an ordered core spanning residues V306-F378. Comprising essentially the same amino acid sequence as the AD and CTE folds, but different from them in structure, this new fold is compatible with the presence of both 3R and 4R tau isoforms.

Seeding PAD12-ΔG389-I392 recombinant tau with seeds extracted from the frontal cortex of the individual with the ΔG389-I392 mutation did not generate filaments with the Pick fold. Instead, this assembly reaction yielded paired helical filaments (PHFs) with the Alzheimer tau fold. This may have been caused by the presence of some 4R tau-immuno-reactive structures, likely to be PHFs, in the frontal cortex (Figure S5).

## Discussion

The findings presented here demonstrate that filaments extracted from the brains of D252V and ΔG389-I392 tau mutation carriers had the Pick fold, while missense mutations G272V and S320F gave rise to structurally distinct but closely related Pick fold variants. All brain-derived filaments consisted of a single protofilament.

D252V is located at the N-terminus of the ordered core and ΔG389-I392 lies outside it; neither mutation perturbs the core filament structure. An additional density near V252 and the ^364^PGGG^367^ motif in the D252V structure may promote filament formation. By contrast, G272V (within the ^269^QPGGG^273^ motif linking the stem to the long arm of the Pick fold) induced local conformational changes, resulting in a more open filament fold. A related fold with a different conformation of the linking ^269^QPGGG^273^ motif was found in the tau filaments with the S320F mutation at the opposite end of the long arm. Even though this mutation at a distant site could affect the conformation of the linking motif allosterically, it seems more likely that the observed conformation was induced locally by other factors, such as non-proteinaceous cofactors and/or post-translational modifications. Despite variations in the relative orientations between the long arm and the stem of these structures, most cross-β packing interactions were maintained between the canonical Pick fold (15) and the G272V and S320F folds (Figure S6). We therefore classify the G272V and S320F folds as variants of the Pick fold.

In all cases of FTDP-17T studied here, Pick bodies were present in nerve cells (16,20-22). It is not clear why *MAPT* mutations D252V, G272V, S320F and ΔG389-I392 cause the formation of 3R tau filaments with the Pick fold. Previously, we showed that *MAPT* mutation ΔK281 in exon 10 gives rise to the Pick fold (7), probably as the result of the relative over-expression of wildtype 3R tau (25) and its assembly into filaments. For D252V, G272V and S320F, both wild-type and mutant tau proteins were present in the filaments.

Missense mutations G272V and S320F have been shown to reduce the ability of 3R and 4R recombinant tau to promote microtubule assembly (24,26) and it has been suggested that S320F tau allosterically disrupts the protection of motif ^306^VQIVYK^311^ from assembly, resulting in the exposure of this amyloidogenic motif (27). Residues ^306^VQIVYK^311^ are necessary for the assembly of tau into filaments (28,29). A primary effect on alternative mRNA splicing of exon 10 cannot be excluded for mutation ΔG389-I392. At the protein level, removal of I392 may interfere with the interaction of residues ^392^IVYK^395^ with ^306^VQIVYK^311^ (14). Removal of ^392^IVYK^395^ increases the propensity of filament formation by PAD12 tau. Where studied, nerve cells of the human brain expressed similar levels of 3R and 4R tau (30). However, we cannot exclude the possibility that some nerve cells express predominantly 3R tau and that the first D252V, G272V and S320F Pick folds form in those cells. Once a seed has formed, it may be maintained through subsequent seeded aggregation, even in the presence of similar levels of 3R+4R tau.

Knowledge of the structures of tau filaments from the *MAPT* mutation cases that were characterised here may facilitate the development of *in vitro* reconstitution systems for filaments with the Pick fold. This may in turn yield insights into the mechanisms of tau filament assembly (13) and the availability of purified seeds with disease-relevant structures may aid mechanistic studies of seeded aggregation in cells and in animals, as well as the development of structure-specific binders for diagnostic and therapeutic purposes. Only the Alzheimer and the CTE tau folds have so far been reproduced *in vitro* (12,14).

Our attempt at assembling PAD12 tau into Pick filaments through the addition of tau seeds extracted from the brain of an individual with sporadic Pick’s disease (15) yielded only a few filaments that were suitable for cryo-EM structure determination, suggesting that they were partially unfolded. Whereas tau filaments extracted from the cerebral cortex of an individual with sporadic Pick’s disease (15) or individuals with *MAPT* mutation ΔK281 (7) comprise predominantly singlets (narrow Pick filaments) and a minority of doublets (wide Pick filaments), the *in vitro* seeded assembly of PAD12 tau led exclusively to doublets with the Pick fold and an inter-protofilament packing different from that of the wide Pick filaments.

Seeding of recombinant PAD12-ΔG389-I392 tau with seeds from the frontal cortex of the case with the ΔG389-I392 mutation led to the formation of PHFs with the Alzheimer tau fold, probably because of the presence of PHF seeds. It remains unclear why this deletion leads to the formation of Pick filaments in the human brain.

In contrast, seeding of PAD12 recombinant tau with mutations D252V, G272V or S320F led to the formation of abundant singlets and doublets with the Pick tau fold. However, none of those assembly reactions replicated the structures of the seeds exactly. Only the long arm and most of the stem were preserved, whereas alternative conformations were observed for the linking segment ^269^QPGGG^273^ and a small number of residues from both ends of the ordered cores. The stem and the long arm of the Pick fold could be two separate domains with more rigid structures than those of the linking and terminal regions, the conformations of which might differ depending on the conditions of nucleation-dependent assembly. It is possible that seeded assembly with PAD12 tau plus a combination of the D252V, G272V or S320F mutations could lead to filaments that more closely resemble those extracted from human brains, including under nucleation-dependent assembly conditions.

PAD12, as well as its D252V, G272V and S320F mutants formed a doublet that has not been observed in tau filaments extracted from human brains. Since this doublet was stabilised by salt bridges, reconstitution in buffers with higher salt concentrations could perhaps lead to a greater proportion of singlets. We note that other salt-bridged doublets are also commonly found in tau filaments extracted from human brains. The apparent lack of electrostatically-stabilised doublets of the Pick fold in these FTDP-17T mutants may have been the result of post-translational modifications of some residues. In addition to filaments with the Pick fold, PAD12-S320F tau also formed filaments with a fold that has not been observed in human brain samples; its presence may confound studies with these filaments. Surprisingly, the 20° rotation of the long arm relative to the stem of the Pick fold that we observed in the brain-derived S320F filaments was not present in the PAD12-S320F filaments, which adopted a conformation identical to that of the wildtype Pick fold, except for the inclusion of the phenylalanine side chain at the tip of the ordered core. The observation that S320F tau can adopt both orientations raises the possibility that this may also be the case for wildtype tau. To date, the cryo-EM structures of tau filaments from only one case of sporadic Pick’s disease and two cases with *MAPT* mutation ΔK281 have been determined (7,15).

Here we establish that tau filaments extracted from the brains of D252V and ΔG389-I392 mutation carriers have the Pick fold, with mutations G272V and S320F giving rise to variants of the Pick fold. These cryo-EM structures, together with those determined previously (7-11), provide a structure-based classification of cases of FTDP-17T (Figure 8). Tau filaments from cases of FTDP-17T that have the same structures as those from sporadic tauopathies or variants thereof consist of both mutant and wild-type tau (7-9). By contrast, filaments from human cases with *MAPT* mutations that encode P301L or P301T tau have structures unlike those from sporadic tauopathies and are only made of mutant tau (10). Specific tau folds define different molecular pathologies, but the same fold can form in different clinical contexts. Thus, the Alzheimer tau fold is found in AD and in some cases of FTDP-17T (9,31). By contrast, the Pick tau fold (15) is diagnostic of cases of frontotemporal dementias, whether sporadic or inherited. This work provides a foundation for the development of structure-based diagnostic and therapeutic approaches for Pick’s disease.

**Figure 8.**
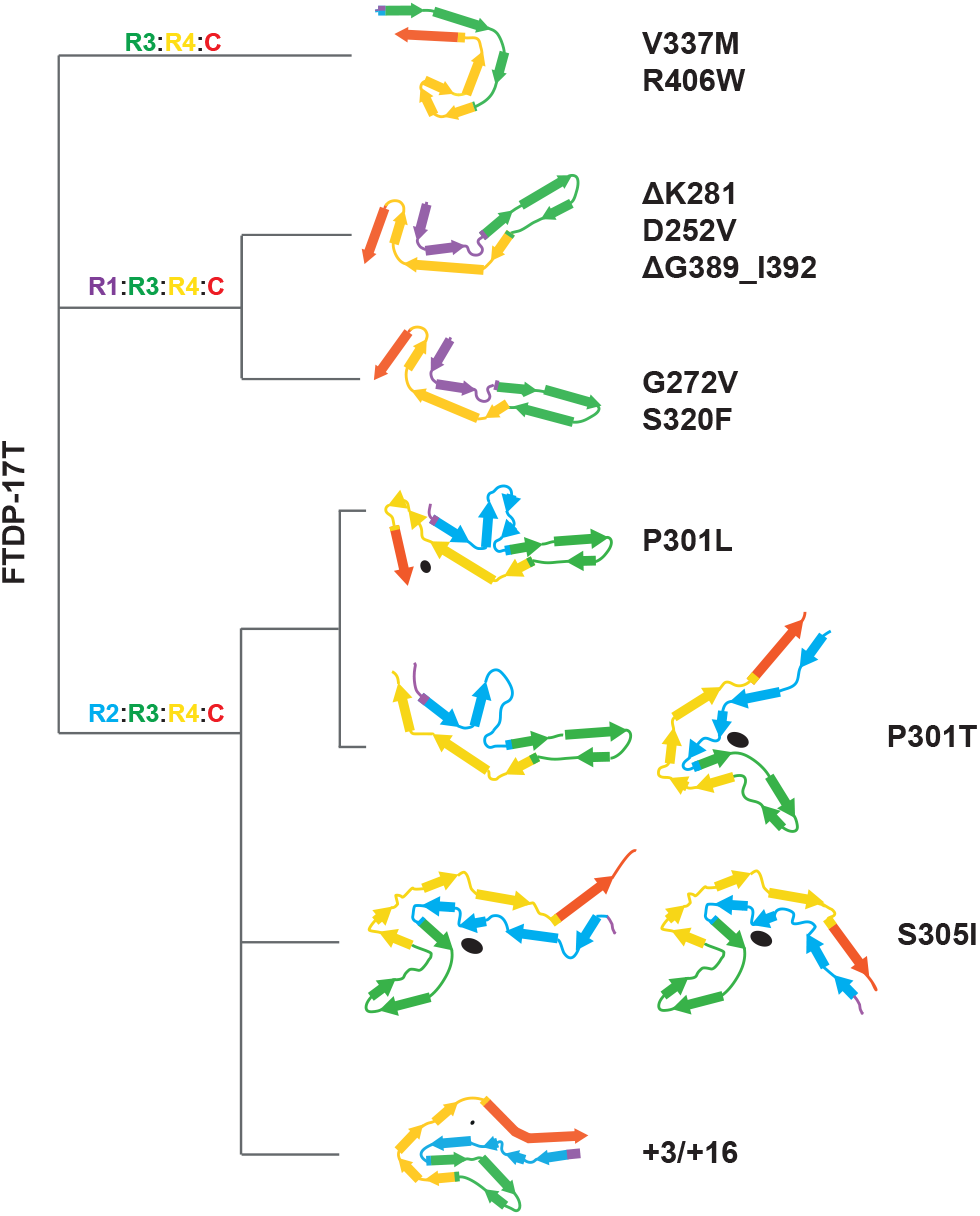
Structure-based classification of cases of FTDP-17T. Dendrogram of inherited tauopathies based on cryo-EM filament structures. Filaments from the brains of individuals with *MAPT* mutations V337M and R406W adopt the Alzheimer tau fold. Filaments from the brains of individuals with *MAPT* mutations D252V and DG389-I392 adopt the Pick tau fold, whereas those with *MAPT* mutations G272V and S320F adopt variants of the Pick fold. Mutations P301L, P301T and S305I in *MAPT* give rise to tau filaments with cryo-EM structures unlike those found in sporadic tauopathies. Mutations +3 and +16 in intron 10 of *MAPT* give rise to the AGD tau fold. Internal, non-proteinaceous densities are indicated in black.

## Data availability

Cryo-EM maps were deposited to the EM Data Bank under the following accession codes: EMD-56597, EMD-56600, EMD-56599 and EMD-56601. Corresponding refined atomic models were deposited to the PDB under the following accession codes: 28LJ, 28LP, 28LO and 28LQ. Please address requests for materials to the corresponding authors.

## Acknowledgements

We thank the participants’ families for donating brain tissues, T. Darling, I. Clayson and J. Grimmett for help with high-performance computing and the EM facility of the Medical Research Council (MRC) Laboratory of Molecular Biology for help with cryo-EM data acquisition. This work was supported by the MRC, as part of UK Research and Innovation (MC_UP_A025_1013 to S.H.W.S. and MC_1051284291 to M.G.). It was also supported by grants from the National Natural Science Foundation of China (32571423 to C.Q.).

## Author contributions

T.T.W., H.S., P.W.C., Z.J. and J.C.v.S. identified participants and performed neuropathology. C.Q. performed immunoblot analysis. C.Q. and S.P.-C carried out mass spectrometry. S.L. and J.S. performed seeded aggregation. C.Q. and S.L. collected cryo-EM data. C.Q., S.L., A.G.M. and S.H.W.S. analysed cryo-EM data. S.H.W.S. and M.G. supervised the project. All authors contributed to the writing of the manuscript.

## Competing interest statement

S.L., M.G. and S.H.W.S. are inventors on a patent application on the generation of disease-specific tau filament structures from PAD12 tau. The other authors declare no competing interests.

## Copyright statement

For the purpose of open access, the MRC Laboratory of Molecular Biology has applied a CC BY public copyright licence to any Author Accepted Manuscript version arising.

## Materials and Methods

### Filament extraction

Sarkosyl-insoluble material was extracted from *postmortem* brain tissues of individuals with mutations encoding D252V, ΔG389_I392, G272V and S320F in *MAPT*, as described (32). Frontal cortex from the case of sporadic Pick’s disease that was previously used for determination of the Pick fold was also used (15). The tissues were homogenised in 20 volumes (w/v) buffer A (10mM Tris-HCl, pH 7.5, 0.8 M NaCl, 10% sucrose, 1mM EGTA). Following homogenisation, 2% sarkosyl was added and the samples were incubated at 37°C for 30 min, followed by a 10 min centrifugation at 7,000g. The resulting supernatants were clarified by ultracentrifugation at 100,000g for 60 min and the pellets were resuspended in buffer A (1 ml/g tissue), followed by a 10 min centrifugation at 9,500 g. The resulting supernatants were diluted 3-fold with buffer B (50mM Tris-HCl, pH 7.5, 0.15M NaCl, 10% sucrose, 0.2% sarkosyl), followed by ultracentrifugation at 100,000 g for 60 min. The final pellets were resuspended in 100 μl/g of buffer C (20mM Tris HCl, pH 7.4, 100mM NaCl) for cryo-EM analysis.

### Immunohistochemistry

Postmortem brain tissue from the ΔG389-I392 *MAPT* mutation case was processed, as described (16). For tau immunohistochemistry, 7 μm thick sections from formalin-fixed and paraffin-embedded anterior frontal cortex tissue were stained with haematoxylin/eosin and immunolabelled for tau phosphorylated at S202 and T205 (AT8, 1:600, Thermo Fisher), 3R tau (8E6/C11, 1:800, Millipore) and 4R tau (1E1/A6, 1:4,000, Millipore) using an automated platform (A. Menarini Diagnostics) with diaminobenzidine as the chromogen and with appropriate controls. Tissue storage was under a Human Tissue Authority licence; appropriate consent and ethical approval from the National Hospital for Neurology and Neurosurgery Research Ethics Committee were in place.

### Immunoblotting

Samples were resolved on 4-20% Bis-Tris gels (NuPage) and the primary antibodies (BR133, 1:1,000; BR134, 1:1,000) were diluted in PBS plus 0.1% Tween20 and 5% non-fat dry milk. Immunoblotting was carried out as described (10).

### Expression and purification of recombinant tau

PAD12 tau with the D252V, G272V, S320F or ΔG389-I392 mutation was produced using in vivo assembly (33) in E. coli BL21(DE3)-gold cells (Agilent Technologies), as described (12). Cells were resuspended in 1 L of 2×TY (tryptone yeast) supplemented with 2.5 mM MgSO4 and 100 mg/L ampicillin and grown to an optical density of 0.8 at 600 nm. They were induced by the addition of 0.6 mM IPTG for 4 h at 37 °C, collected by centrifugation (4,000g for 30 min at 4 °C) and flash frozen. The pellets were resuspended in washing buffer at room temperature (50 mM MES pH 6.5, 250 mM NaCl, 10 mM EDTA, 10 mM DTT, supplemented with 0.03 mM chymostatin, 0.1 mM phenylmethylsulphonyl fluoride (PMSF), 0.1 mM 4-(2-aminoethyl)benzenesulphonyl fluoride hydrochloride (AEBSF), supplemented with cOmplete EDTA-free protease inhibitor cocktail (Roche) (three tablets per litre and an additional tablet in 50 ml, 40 mg/ml DNAse I (Sigma) and 10 mg/ml bovine pancreas RNase (Sigma) were added). Cells were lysed by sonication (90% amplitude using a Sonics VCX-750 Vibracell ultrasonic processor for 4 min with 3s on and 6 s off) at 4°C. The lysed cells were centrifuged at 20,000g for 35 min at 4 °C, and the lysates diluted 5-fold to reach a NaCl concentration of 50 mM, followed by loading onto a HiTrap CaptoS 5-ml column (GE Healthcare). The column was washed with ten volumes of buffer A with 50 mM NaCl, followed by elution through a gradient of buffer A containing 0–1 M NaCl. Fractions of 3.5 ml were collected and analysed by SDS–PAGE (4–20% Tris–glycine gels). Protein-containing fractions were pooled and precipitated using 0.38 g/ml ammonium sulphate and left on a rocker for 1h at 4 °C. The solution was then centrifuged at 20,000g for 35 min at 4 °C and resuspended in 2 ml of 10 mM potassium phosphate buffer pH 7.2 containing 10 mM DTT and loaded onto a 16/600 75-pg size-exclusion column. Fractions were analysed by SDS–PAGE and protein-containing fractions pooled and concentrated at 4 °C to 20 mg/ml using molecular weight concentrators with a cutoff filter of 3 kDa. Purified protein samples were flash-frozen in 50–100 µl aliquots. Protein concentrations were determined using a NanoDrop 2000 (Thermo Fisher Scientific).

### Seeded assembly of recombinant PAD12 tau

Prior to assembly, proteins and buffers were filtered (Costar Spin X centrifuge tube filters, 0.22 mm), followed by determination of the protein concentrations. Assembly reactions were prepared in Eppendorf protein LoBind tubes. Reactions were prepared at room temperature by mixing water, buffering agent (from a 1M HEPES stock, pH 7.28), TCEP (from a 100 mM stock), salt (from a 1M sodium citrate) and thioflavin T (from a 150 mM stock). The concentrations were: 50 mM protein, 40 mM HEPES, 200 mM sodium citrate, 4mM TCEP, 3 mM thioflavin T. Filaments from postmortem brains of individuals with *MAPT* mutations were extracted as described above. Sarkosyl-insoluble pellets were diluted 100-fold (100,000 ml/g tissue) and 1 ml was used per seeding reaction in a 30 ml reaction volume (0.33 mg/ml) and added prior to the addition of thioflavin T. Prior to reaction setup, wells in a 384-well plate were rinsed with 100 ml water. Reactions were prepared in batch and 30 ml aliquots were dispensed into each well. Each reaction had an empty well next to it, to prevent cross-contamination. Plates were incubated at 37° C with orbital shaking (5s on, 5s off, 500 rpm) and thioflavin T fluorescence was measured every 10 min.

### Mass spectrometry

Mass spectrometry was carried out as described (9). In brief, sarkosyl-insoluble pellets were resuspended in 200 µl of hexafluoroisopropanol. Following a 3-min sonication at 50% amplitude (QSonica), they were incubated at 37 °C for 2 h and centrifuged at 100,000g for 15 min, before being dried by vacuum centrifugation. Protein samples that were resuspended in 4 M urea and 50 mM ammonium bicarbonate were reduced with 5 mM DTT at 37 °C for 40 min and alkylated with 10 mM chloroacetamide for 30 min. After dilution to 1 M urea, the G272V sample was digested with LysC (Promega), the D252V sample with trypsin (Promega), the S320F sample with chymotrypsin (Promega) and the ΔG389_I392 sample with AspN (Promega). All samples were incubated overnight at 25 °C. Digestion was stopped by the addition of formic acid to a final concentration of 0.5%, followed by centrifugation at 16,000g for 5 min. The supernatants were desalted using homemade C18 stage tips (3 M Empore) packed with Poros Oligo R3 (Thermo Fisher Scientific) resin.

Bound peptides were eluted stepwise with 30-80% acetonitrile and partially dried in a SpeedVac concentrator (Savant). Samples were analysed by liquid chromatography (LC)–MS/MS using a Orbitrap Eclipse Tribrid mass spectrometer (Thermo Fisher Scientific) coupled online to a Vanquish Neo Nano LC system (Thermo Fisher Scientific). LC–MS/MS data were searched against the UP000005640_9606_human_proteome (UniProt, downloaded 2023), supplemented with Tau mutated sequences, using the Mascot search engine (Matrix Science, version 2.80). Scaffold (version4, Proteome Software) was used to validate MSMS-based peptide and protein identifications.

### Cryo-EM sample preparation and data collection

Prior to freezing, the extracted filaments sample was centrifuged at 3,000g for 1 min. The cryo-EM grids (Quantifoil 1.2/1.3, 300mesh) were glow-discharged for 1 min using an Edwards (S150B) sputter coater. A 3μl aliquot of the sample was applied to each glow-discharged grid. The grid was blotted with filter paper (blot force 10, wait time 6 seconds, blot time 4s) and plunge-frozen into liquid ethane using Vitrobot MarkIV (FEI) (100% humidity and 4°C). Cryo-EM images were collected on Titan Krios electron microscopes (FEI, 300kV) equipped with a Falcon4i or K3 direct electron detector. Images were recorded with a total dose of 40 e/Å^2^ and a pixel size of 0.824 Å or 0.744 Å for Falcon4i detector (or 0.826 Å for K3 detector). See Tables S1 and S2 for further details.

### Cryo-EM data processing

All data processing was performed using RELION software package (34-37). The cryo-EM images were corrected with gain reference. Motion-correction and dose weighting were performed using RELION’s own implementation. Contrast transfer function (CTF) was estimated using CTFFIND-4.1 (38). Filaments were manually picked, and segments were extracted with a box size of 1024 pixels and downscaled to 256 pixels. Reference-free 2D classification was performed to remove poor-quality particles. Selected class averages were re-extracted using a box size of 400 pixels. *relion_helix_inimodel2d* was used to generate initial models *de novo* from 2D class averages based on filament crossover distances. 3D refinement was then performed in RELION, with further optimization of the helical twist and rise using local searches. For the S320F dataset, 3D classification was performed using Blush regularisation (39). The selected good class was further refined using amyloid Blush regularisation. To further improve the resolution, Bayesian polishing and CTF refinement were performed (40,41). Final maps were sharpened using post-processing procedures in RELION, and resolution was determined based on gold-standard Fourier shell correlations (FSC) at the 0.143 criterion (42). *relion_helix_toolbox* was used to impose helical symmetry on the post-processing maps (Figures S7-S10).

### Model building and refinement

Atomic models were built manually using Coot (43), using previously published structures as references (15,31,44). Model refinements were performed using either *Servalcat* or ISOLDE (45,46) and REFMAC5 (47,48). Models were validated with MolProbity (49). Figures were prepared with ChimeraX (50) and PyMOL (51).

---

**Figure S1.**
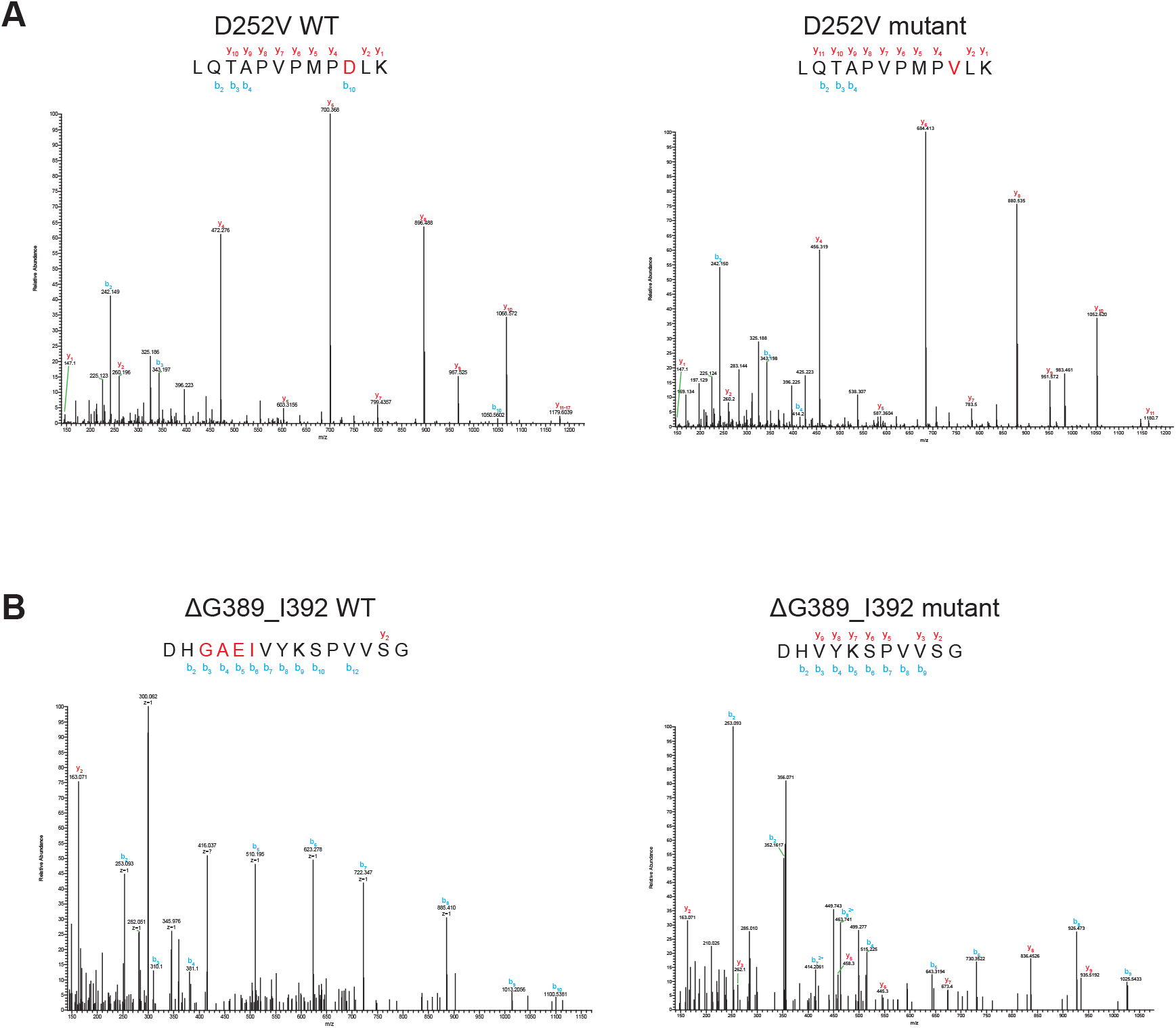
Mass spectrometric analysis of tau from the sarkosyl-insoluble fractions of cases with *MAPT* mutations encoding D252V tau (frontal cortex) and ΔG389_I392 tau (frontal cortex). **A**, MSMS mass spectra of sarkosyl-insoluble tau from the D252V case. **B**, MALDI mass spectrum of sarkosyl-insoluble tau from the ΔG389_I392 case. In both cases, peptides corresponding to wild-type (WT) and mutant tau were detected.

**Figure S2.**
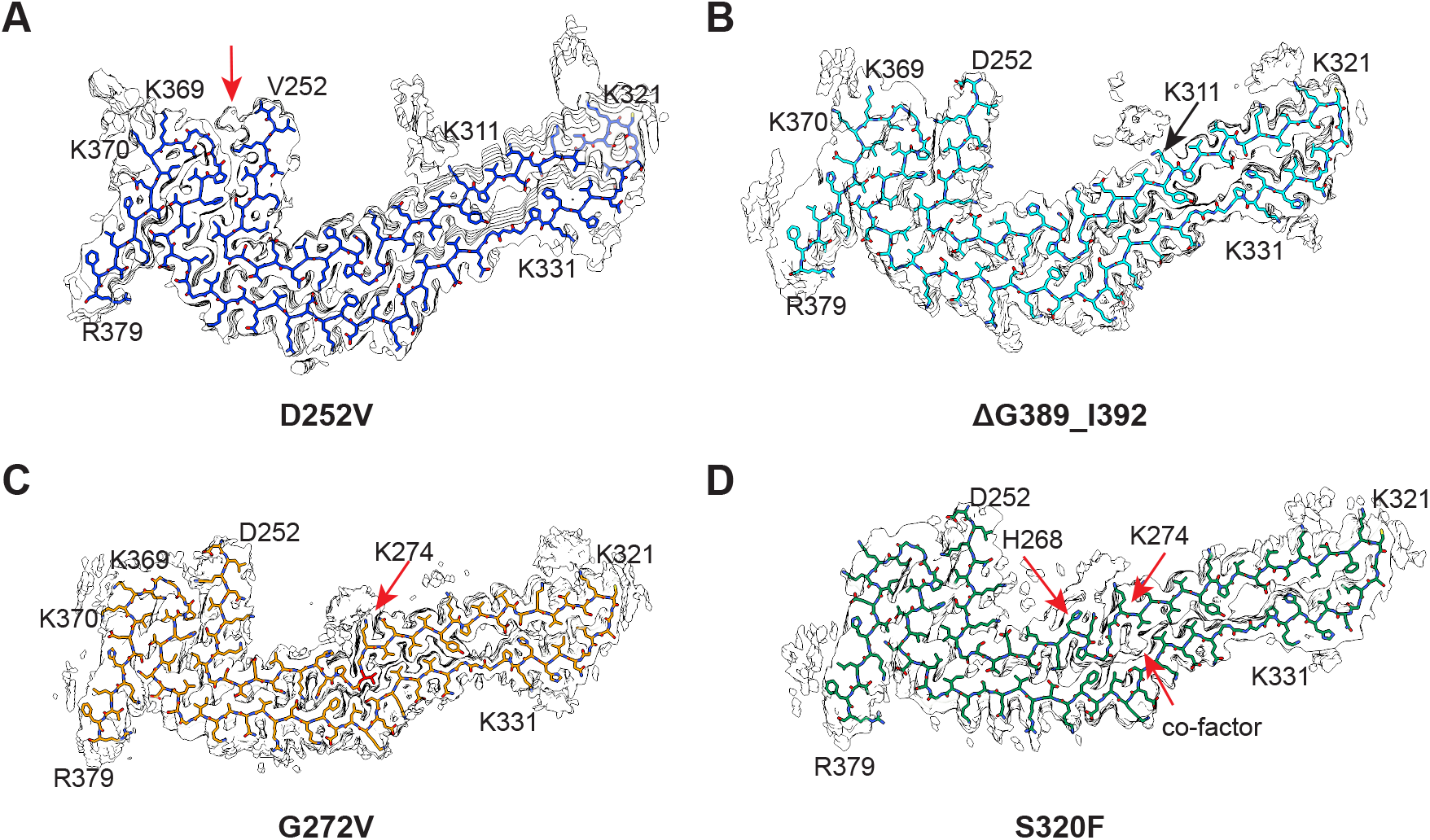
Cryo-EM maps of tau filaments from individuals with *MAPT* mutations D252V, ΔG389_I392, G272V and S320F, showing at low contour levels to reveal additional densities on the filament surface. **A**, Cryo-EM map of D252V tau filaments. The red arrow indicates the cofactor located near residue V252. **B**, Cryo-EM map of ΔG389_I392 tau filaments. **C**, Cryo-EM map of G272V tau filaments. The red arrow indicates the additional density near residue K274. **D**, Cryo-EM map of S320F tau filaments, the red arrows indicate the cofactor near G272 and the additional density near H268 and K274.

**Figure S3.**
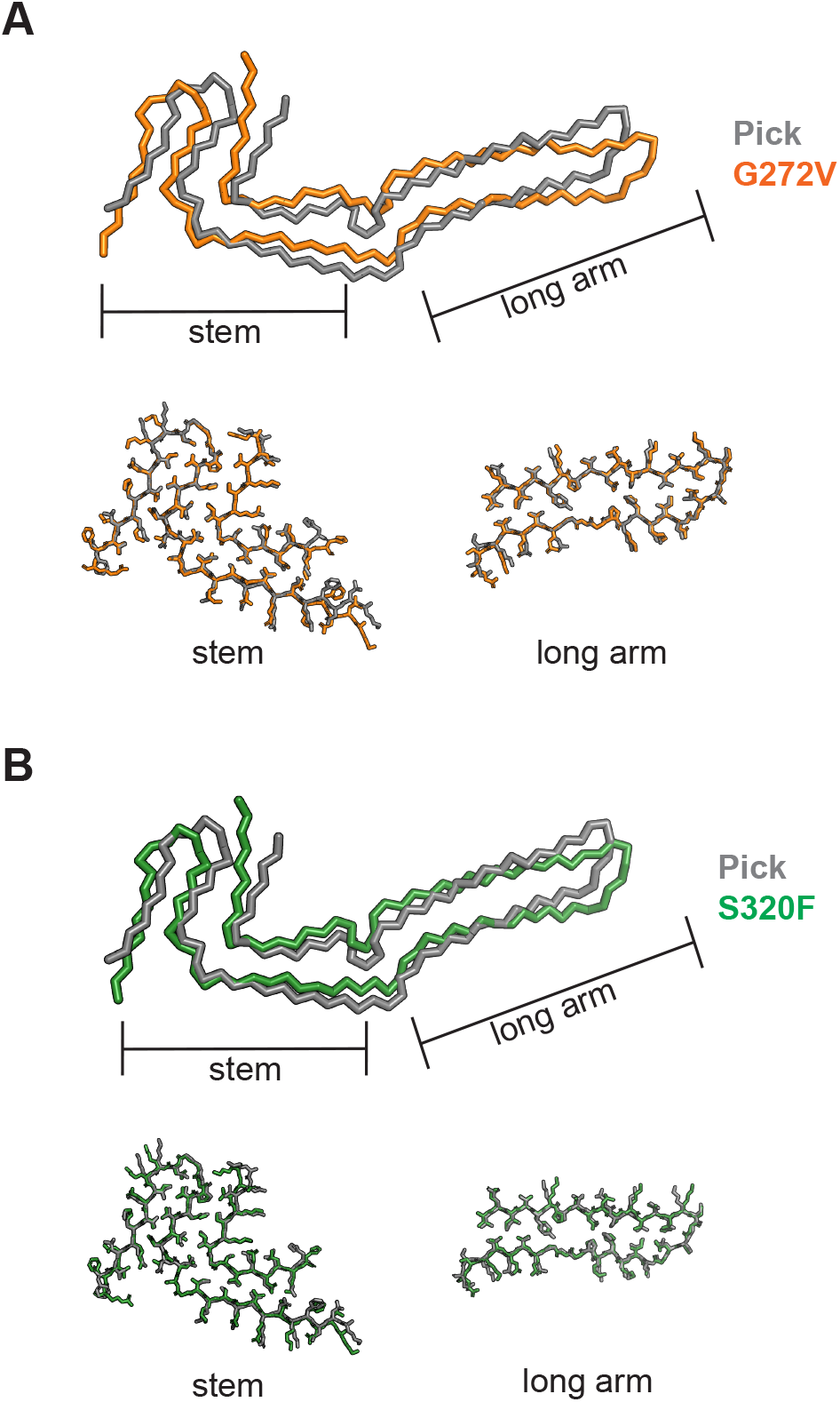
Comparison of G272V and S320F tau filaments with those from sporadic Pick’s disease. **A**, Overlay of structures with the G272V mutation (orange) with the Pick fold (grey). The RMSD between Cα atoms of the two structures is 4.833Å. When the stem region and the long arm region are aligned separately, the RMSD values are 0.408 Å and 0.476 Å. **B**, Overlay of structures with the S320F mutation (green) with the Pick fold (grey). The RMSD between Cα atoms of the two structures is 3.752 Å. When aligning the stem region and the long arm region separately, RMSD values were 0.603 Å and 0.599 Å, respectively.

**Figure S4.**
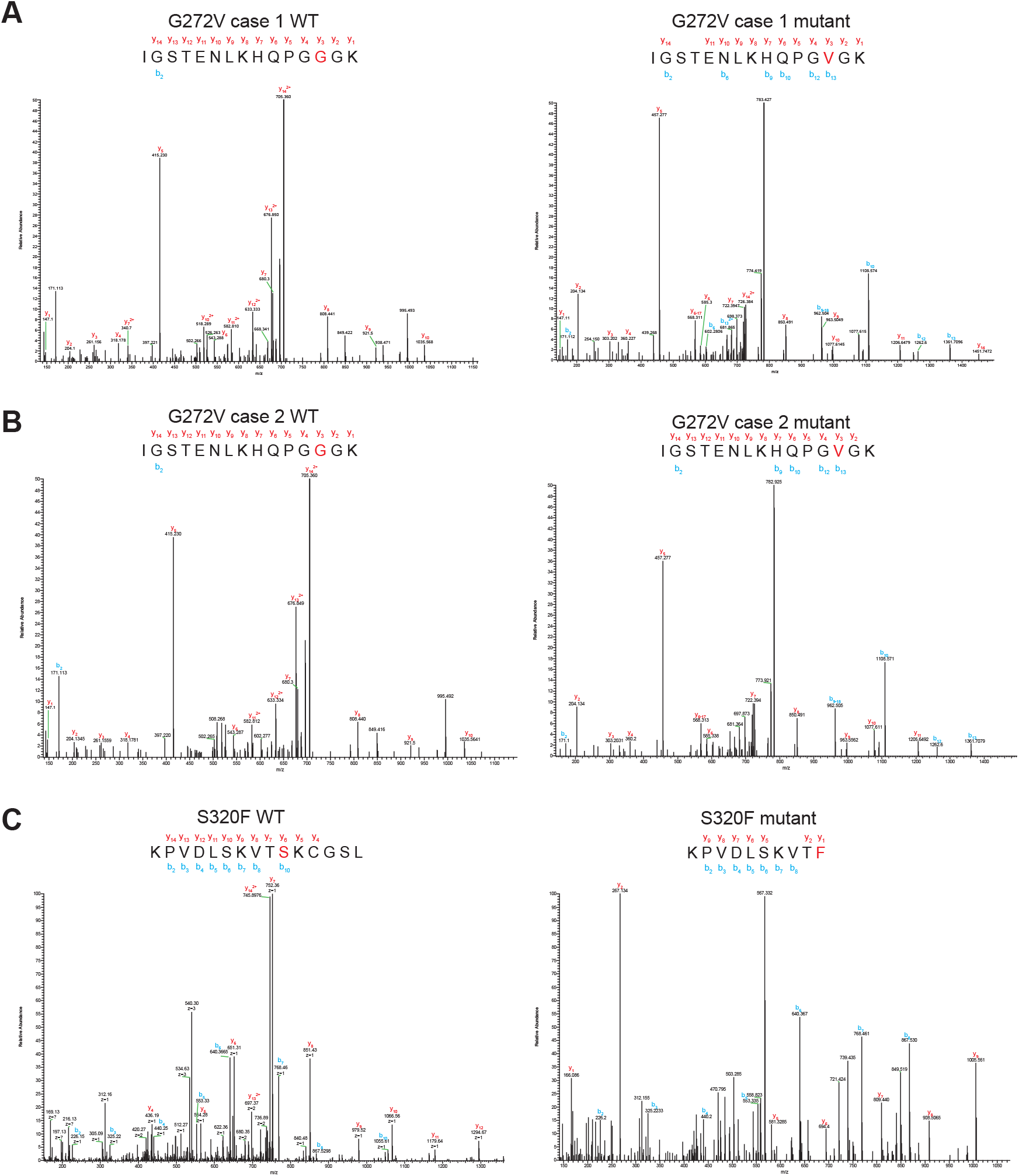
Mass spectrometric analysis of tau from the sarkosyl-insoluble fractions of cases with *MAPT* mutations encoding G272V tau (temporal cortex) and S320F tau (caudate nucleus). **A**, MSMS mass spectra of tau filaments extracted from G272V case 1. **B**, MSMS mass spectra of tau filaments extracted from G272V case 2. **C**, MSMS mass spectra of tau filaments extracted from the S320F case. Peptides corresponding to both wildtype (WT) and mutant (V272) tau were detected.

**Figure S5.**
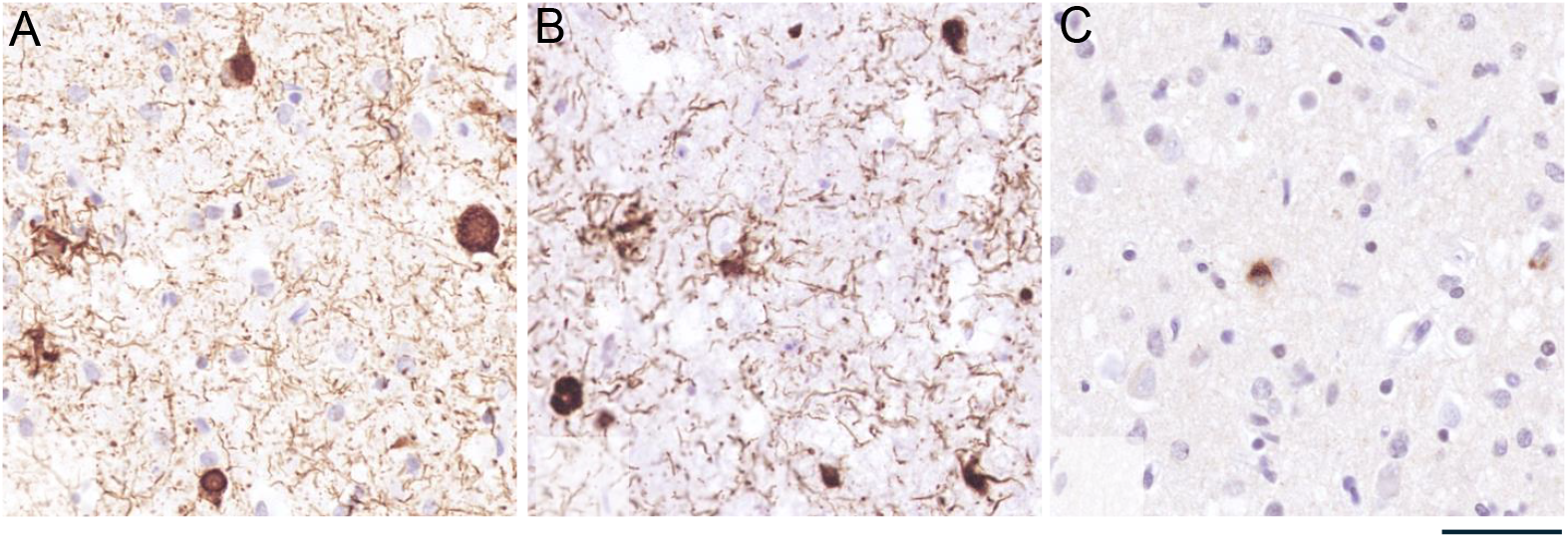
Tau pathology in the frontal cortex of the proband with *MAPT* mutation ΔG389-I392. **A**, AT8 immunohistochemistry shows a dense meshwork of tau-positive neuropil threads, together with abundant neuronal cytoplasmic inclusions showing tangle, pre-tangle and Pick body-shaped morphologies, as well as astrocytic tau pathology consisting of ramified astrocytes. **B**, 3R tau immunostaining shows similar morphologies and distributions as for AT8, confirming the predominance of 3R tau-positive neuronal and glial inclusions. **C**, 4R tau immunostaining shows rare neuronal cytoplasmic inclusions.

**Figure S6.**
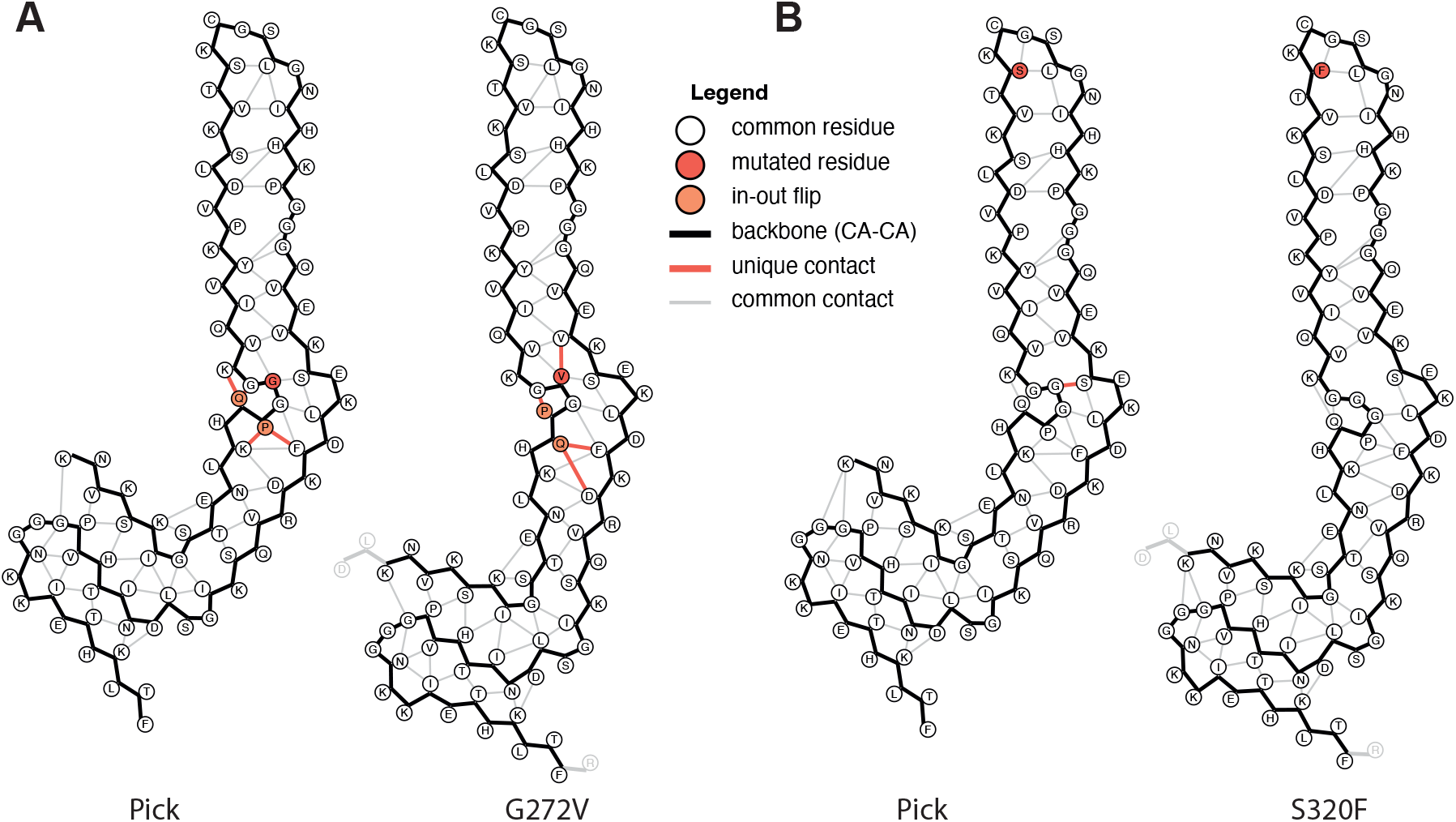
Amyloid packing difference comparisons. **A**, Comparison between the Pick fold and the G272V fold. **B**, Comparison between the Pick fold and the S320F fold. Side chain packing interactions (i.e. residue pairs with any side chain atoms, including Cα atoms for glycines, below 4.5 Å from each other) are shown in grey lines if they exist in both folds and in dark orange lines if they only exist in one of the folds. Mutated residues are highlighted in red circles; residues that have a different side chain orientation, i.e. facing either to the inside or the outside of the ordered core, are highlighted in orange circles. The amyloid packing difference (APD), a new metric for the quantitative comparison of amyloid folds (52), which is calculated as the percentage of residues that are unique in the largest fold that are engaged in distinct unique contacts in one of the folds, or that have a different inside-out orientation between the two folds, is 12% for the Pick-G272V and 5% for the Pick-S320F comparisons.

**Figure S7.**
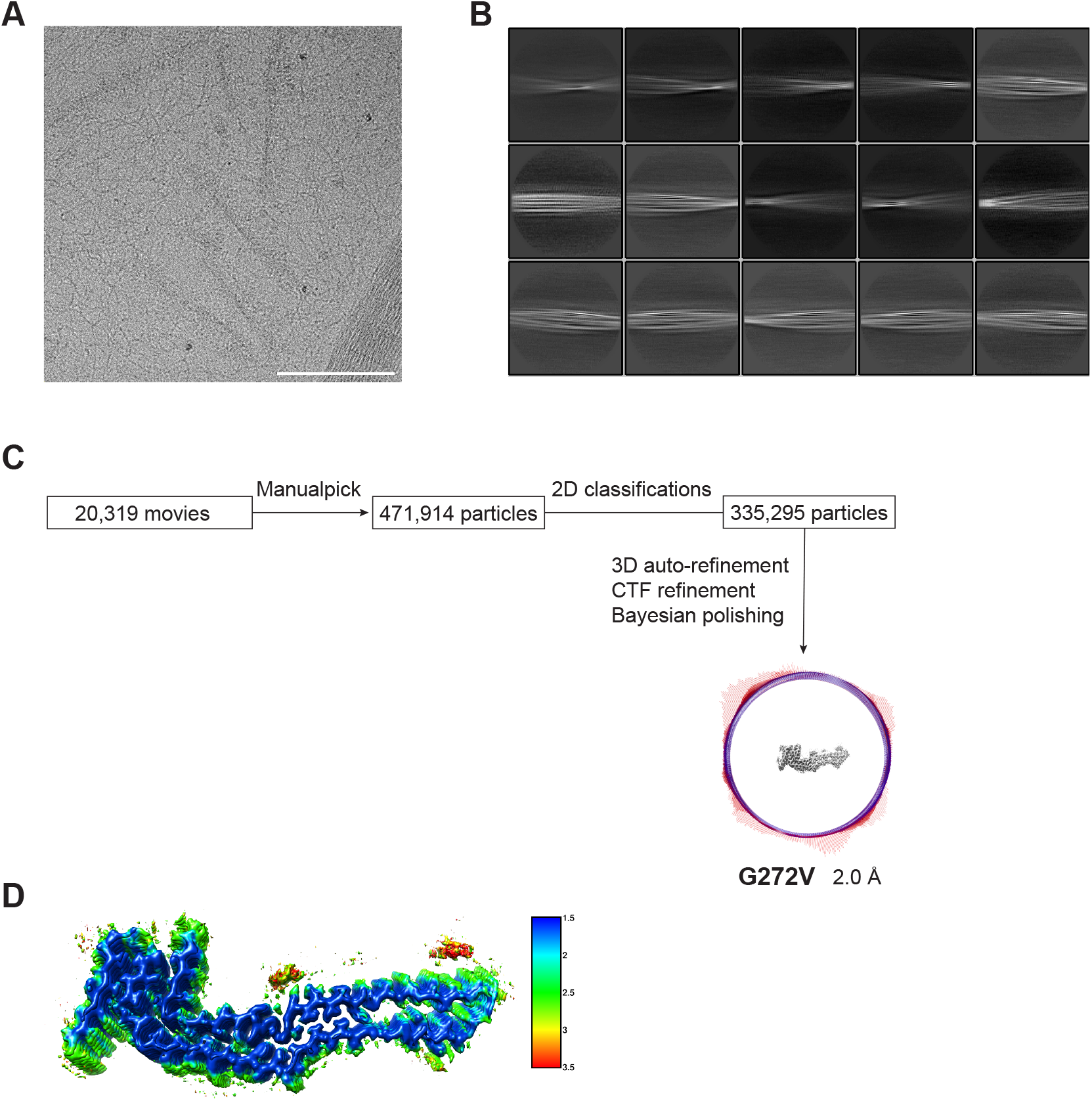
Cryo-EM image processing workflow for the G272V tau filaments. **A**, A representative motion-corrected micrograph; the scale bar, 100 nm. **B**, Representative 2D class averages; the box size is 762 Å. **C**, The overview of the cryo-EM imaging processing workflow. The angular distribution histogram of the final reconstruction is represented at the bottom. **D**, Local resolution maps of G272V tau estimated using RELION.

**Figure S8.**
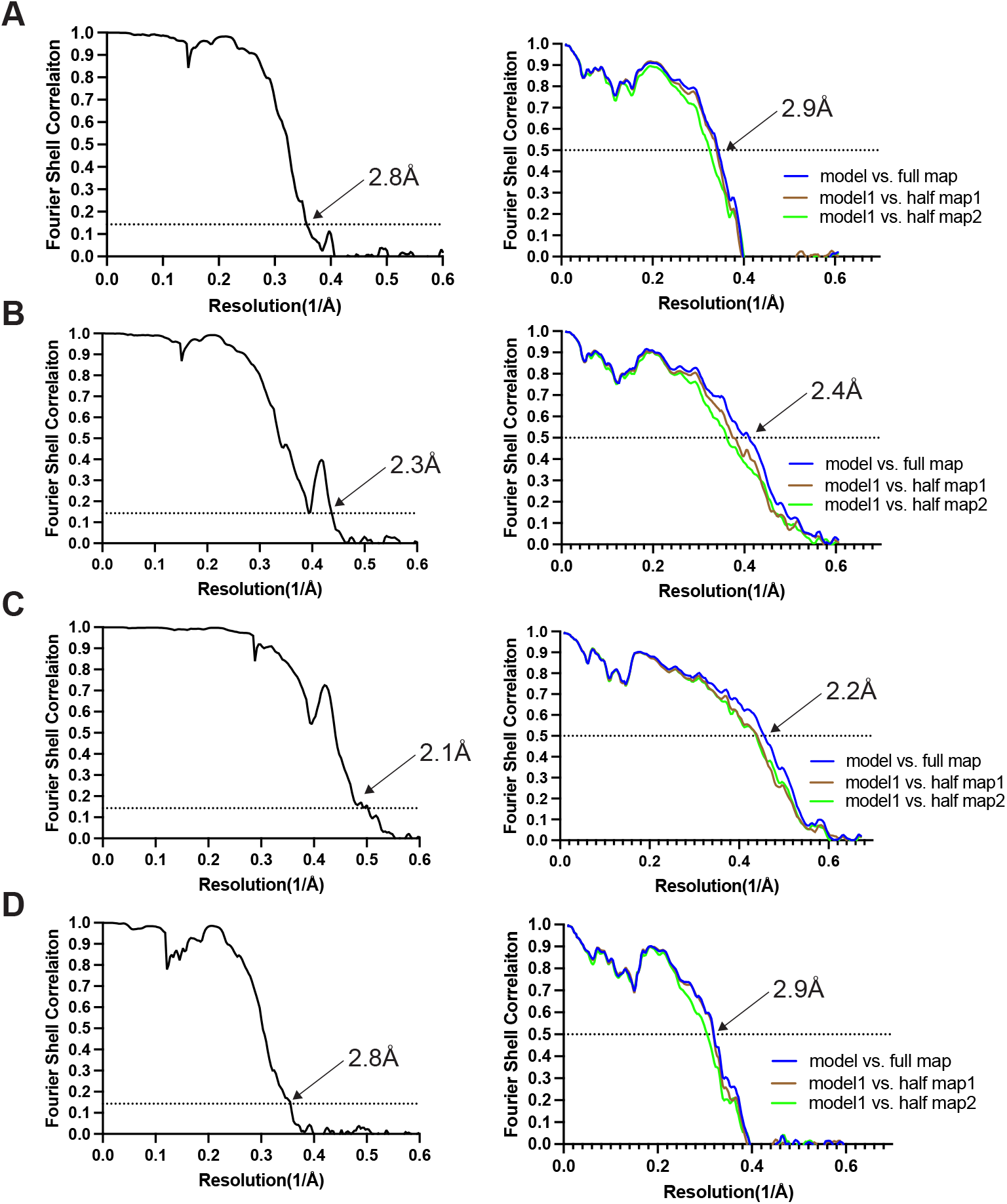
Fourier shell correlation (FSC) curves for cryo-EM maps (left panel) and model to map validation (right panel). **A,** D252V tau structure. **B**, ΔG389_I392 tau structure. **C**, G272V tau structure. **D**, S320F tau structure.

**Figure S9.**
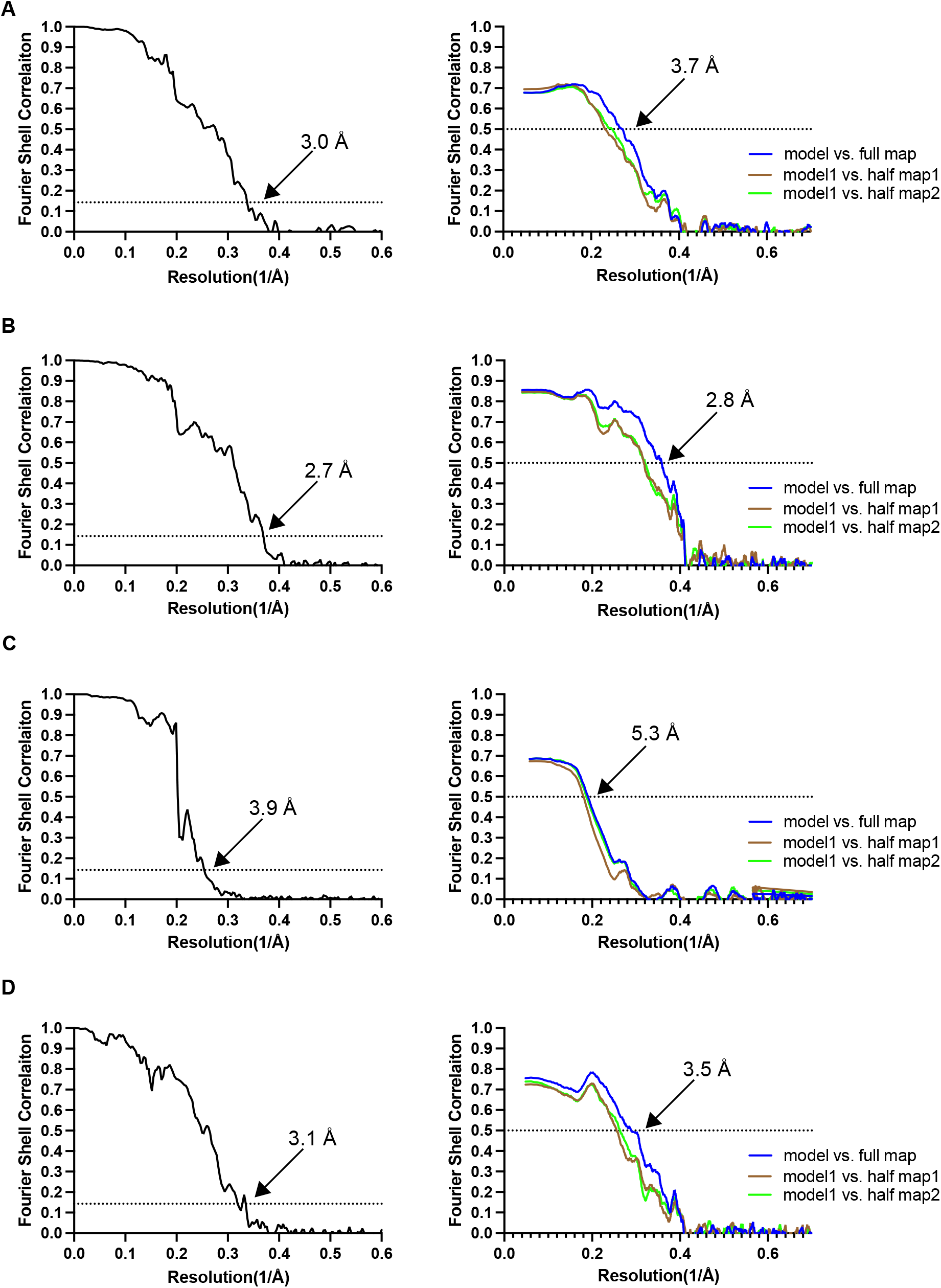
Fourier shell correlation (FSC) curves for cryo-EM maps (left panel) and model to map validation (right panel). **A**, Recombinant D252V tau singlet. **B**, Recombinant D252V tau doublet. **C**, Recombinant G272V tau singlet. **D**, Recombinant G272V tau doublet.

**Figure S10.**
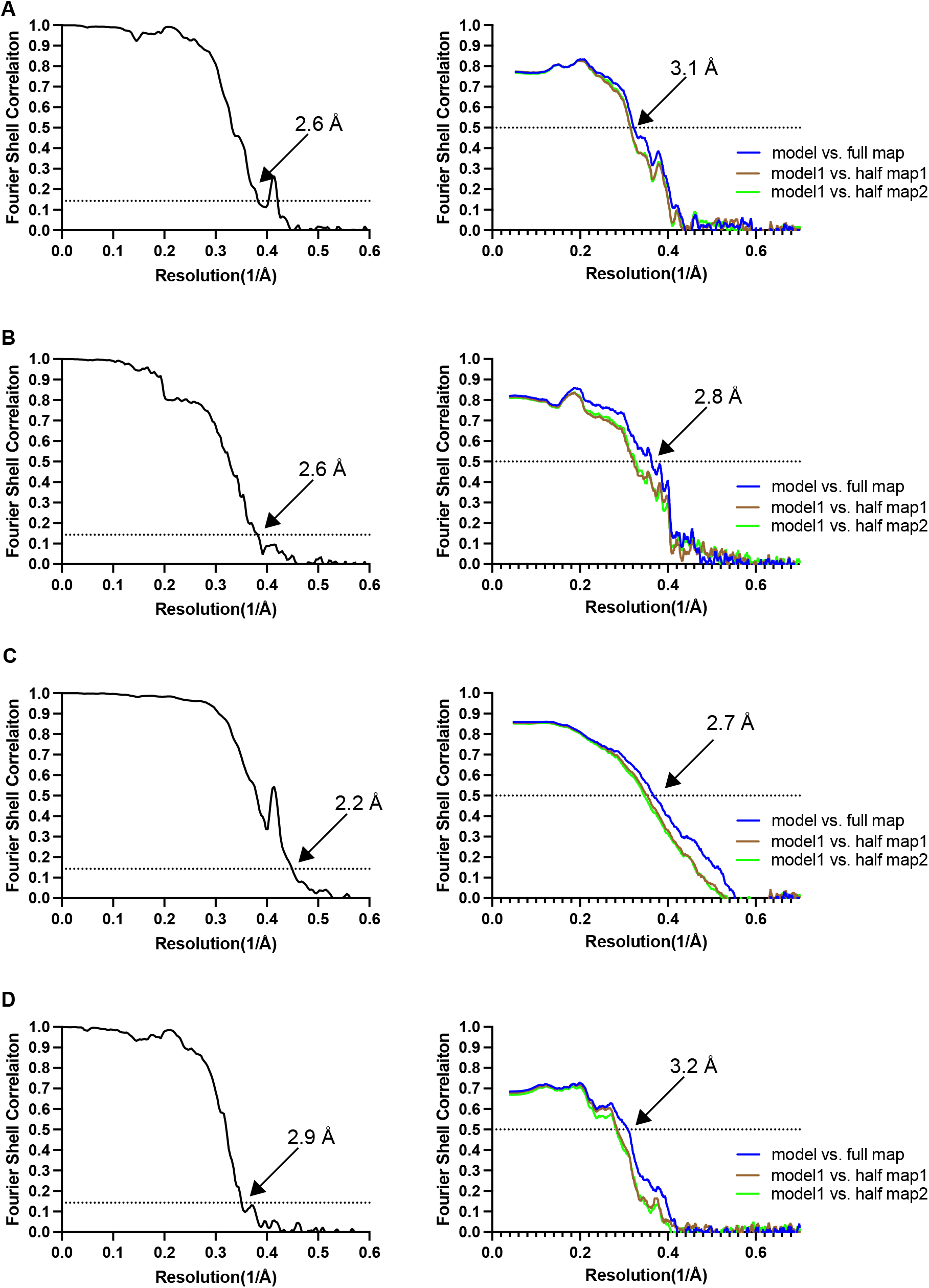
Fourier shell correlation (FSC) curves for cryo-EM maps (left panel) and model to map validation (right panel). **A**, Recombinant S320F tau Pick singlet. **B**, Recombinant S320F tau Pick doublet. **C**, Recombinant S320F tau type 2 singlet. **D**, Recombinant S320F tau type 2 doublet.

**Table S1.** Cryo-EM data collection, refinement and validation statistics.

|  | D252V | ΔG389_I392 | G272V<br>Case 2 | S320F |
| --- | --- | --- | --- | --- |
| **Data collection** |  |  |  |  |
| Microscope | Titan Krios |  |  |  |
| Voltage (kV) | 300 |  |  |  |
| Detector | Falcon4i | K3 | Falcon4i | Falcon4i |
| Electron exposure (e-/Å<sup>2</sup>) | 40 | 40 | 40 | 40 |
| Defocus range (μm) | -1.0 to -2.0 | -1.0 to -2.0 | -1.0 to -2.0 | -1.0 to -2.0 |
| Pixel size (Å) | 0.824 | 0.826 | 0.744 | 0.826 |
| **Data processing** |  |  |  |  |
| Box size (pixel) | 400 | 400 | 400 | 400 |
| Symmetry imposed | C1 | C1 | C1 | C1 |
| Initial particle images (no.) | 196,865 | 321,528 | 471,914 | 331,114 |
| Final particle images (no.) | 124,055 | 143,931 | 335,295 | 40,748 |
| Map resolution (Å)<br>FSC threshold 0.143 | 2.8 | 2.3 | 2.1 | 2.8 |
| Helical rise (Å) | 4.83 | 4.80 | 4.89 | 4.86 |
| Helical twist (°) | -0.7 | -0.73 | -0.66 | -0.65 |
| **Refinement** |  |  |  |  |
| Model resolution (Å)<br>FSC threshold 0.5 | 2.9 | 2.4 | 2.2 | 2.9 |
| Map sharpening *B* factor (Å<sup>2</sup>) | -40 | -34 | -27 | -39 |
| Model composition |  |  |  |  |
| Non-hydrogen atoms | 2948 | 3690 | 2223 | 2972 |
| Protein residues | 388 | 485 | 291 | 388 |
| Ligands | 0 | 0 | 0 | 0 |
| *B* factors (Å<sup>2</sup>) |  |  |  |  |
| Protein | 52.9 | 48.1 | 41.4 | 52.8 |
| R.m.s. deviations |  |  |  |  |
| Bond lengths (Å) | 0.0084 | 0.0048 | 0.0049 | 0.0085 |
| Bond angles (°) | 1.983 | 1.438 | 1.371 | 1.797 |
| Validation |  |  |  |  |
| MolProbity score | 1.88 | 1.87 | 1.57 | 2.65 |
| Clashscore | 3.8 | 5.43 | 1.32 | 8.23 |
| Poor rotamers (%) | 3.57 | 2.38 | 3.53 | 8.33 |
| Ramachandran plot |  |  |  |  |
| Favoured (%) | 95.79 | 95.79 | 95.79 | 91.58 |
| Allowed (%) | 4.21 | 4.21 | 4.21 | 8.42 |
| Disallowed (%) | 0 | 0 | 0 | 0 |
| EMDB | EMD-56597 | EMD-56600 | EMD-56599 | EMD-56601 |
| PDB | 28LJ | 28LP | 28LO | 28LQ |

**Table S2.**
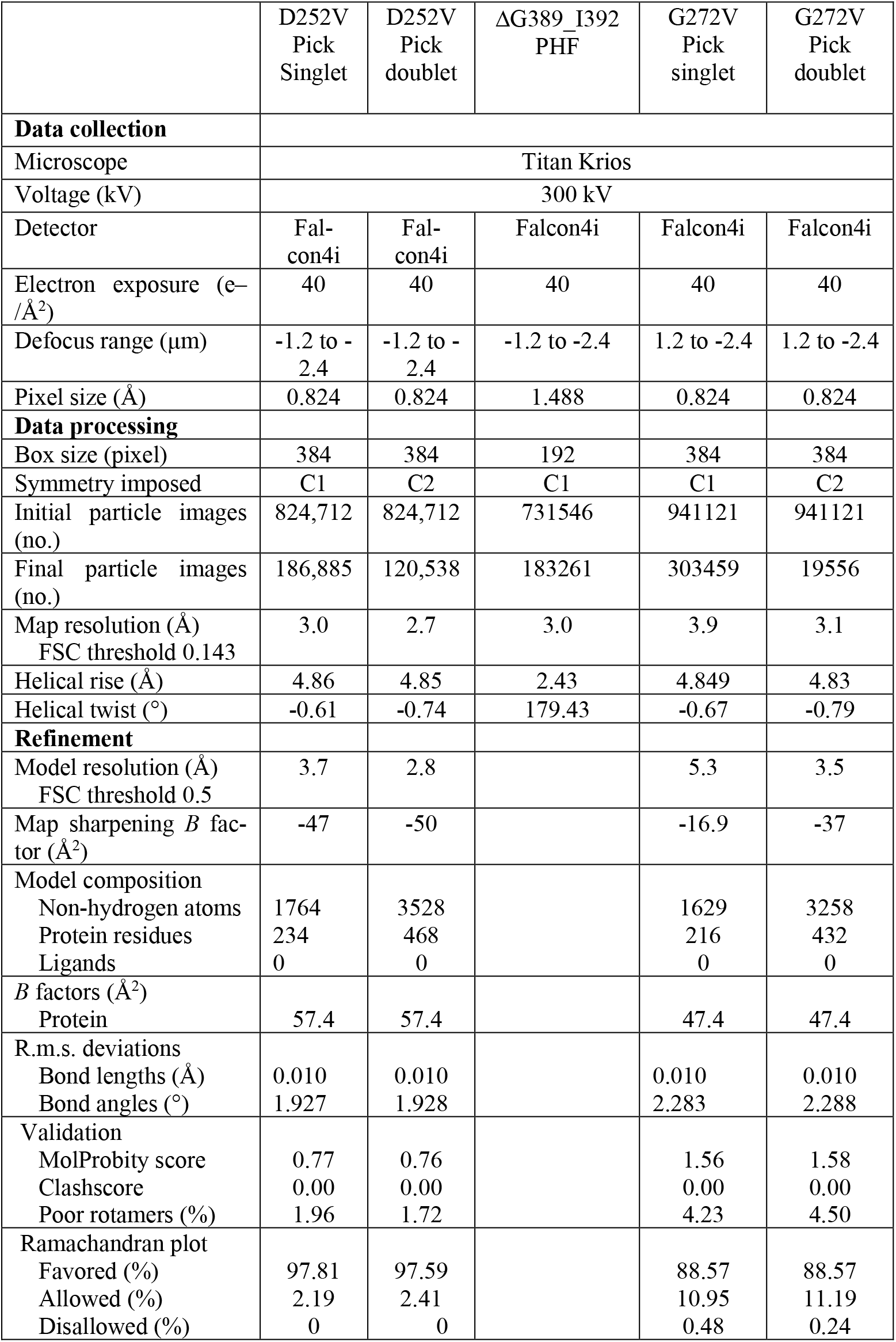
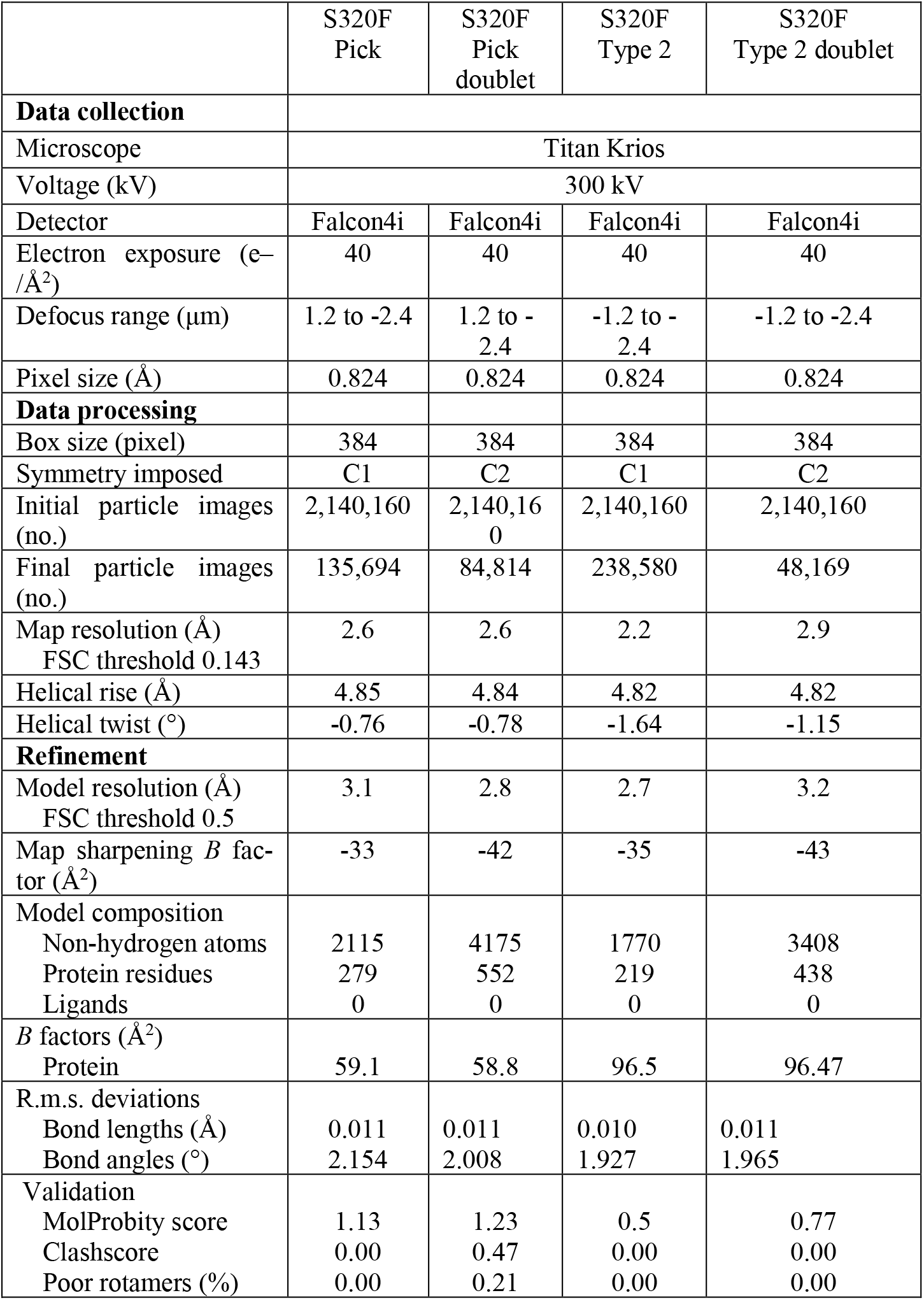
Cryo-EM data collection, refinement and validation statistics.

| | D252V<br>Pick<br>Singlet | D252V<br>Pick<br>doublet | ΔG389_I392<br>PHF | G272V<br>Pick<br>singlet | G272V<br>Pick<br>doublet |
| --- | --- | --- | --- | --- | --- |
| **Data collection** |  |  |  |  |  |
| Microscope | Titan Krios |  |  |  |  |
| Voltage (kV) | 300 kV |  |  |  |  |
| Detector | Fal-<br>con4i | Fal-<br>con4i | Falcon4i | Falcon4i | Falcon4i |
| Electron exposure (e<sup>-</sup>/<br>Å<sup>2</sup>) | 40 | 40 | 40 | 40 | 40 |
| Defocus range (μm) | -1.2 to -<br>2.4 | -1.2 to -<br>2.4 | -1.2 to -2.4 | 1.2 to -2.4 | 1.2 to -2.4 |
| Pixel size (Å) | 0.824 | 0.824 | 1.488 | 0.824 | 0.824 |
| **Data processing** |  |  |  |  |  |
| Box size (pixel) | 384 | 384 | 192 | 384 | 384 |
| Symmetry imposed | C1 | C2 | C1 | C1 | C2 |
| Initial particle images<br>(no.) | 824,712 | 824,712 | 731546 | 941121 | 941121 |
| Final particle images<br>(no.) | 186,885 | 120,538 | 183261 | 303459 | 19556 |
| Map resolution (Å)<br>FSC threshold 0.143 | 3.0 | 2.7 | 3.0 | 3.9 | 3.1 |
| Helical rise (Å) | 4.86 | 4.85 | 2.43 | 4.849 | 4.83 |
| Helical twist (°) | -0.61 | -0.74 | 179.43 | -0.67 | -0.79 |
| **Refinement** |  |  |  |  |  |
| Model resolution (Å)<br>FSC threshold 0.5 | 3.7 | 2.8 |  | 5.3 | 3.5 |
| Map sharpening *B* fac-<br>tor (Å<sup>2</sup>) | -47 | -50 |  | -16.9 | -37 |
| Model composition |  |  |  |  |  |
| Non-hydrogen atoms | 1764 | 3528 |  | 1629 | 3258 |
| Protein residues | 234 | 468 |  | 216 | 432 |
| Ligands | 0 | 0 |  | 0 | 0 |
| *B* factors (Å<sup>2</sup>) |  |  |  |  |  |
| Protein | 57.4 | 57.4 |  | 47.4 | 47.4 |
| R.m.s. deviations |  |  |  |  |  |
| Bond lengths (Å) | 0.010 | 0.010 |  | 0.010 | 0.010 |
| Bond angles (°) | 1.927 | 1.928 |  | 2.283 | 2.288 |
| Validation |  |  |  |  |  |
| MolProbity score | 0.77 | 0.76 |  | 1.56 | 1.58 |
| Clashscore | 0.00 | 0.00 |  | 0.00 | 0.00 |
| Poor rotamers (%) | 1.96 | 1.72 |  | 4.23 | 4.50 |
| Ramachandran plot |  |  |  |  |  |
| Favored (%) | 97.81 | 97.59 |  | 88.57 | 88.57 |
| Allowed (%) | 2.19 | 2.41 |  | 10.95 | 11.19 |
| Disallowed (%) | 0 | 0 |  | 0.48 | 0.24 |

|  | S320F<br>Pick | S320F<br>Pick<br>doublet | S320F<br>Type 2 | S320F<br>Type 2 doublet |
| --- | --- | --- | --- | --- |
| **Data collection** |  |  |  |  |
| Microscope | Titan Krios |  |  |  |
| Voltage (kV) | 300 kV |  |  |  |
| Detector | Falcon4i | Falcon4i | Falcon4i | Falcon4i |
| Electron exposure (e<sup>-</sup>/Å<sup>2</sup>) | 40 | 40 | 40 | 40 |
| Defocus range (µm) | 1.2 to -2.4 | 1.2 to -2.4 | -1.2 to -2.4 | -1.2 to -2.4 |
| Pixel size (Å) | 0.824 | 0.824 | 0.824 | 0.824 |
| **Data processing** |  |  |  |  |
| Box size (pixel) | 384 | 384 | 384 | 384 |
| Symmetry imposed | C1 | C2 | C1 | C2 |
| Initial particle images (no.) | 2,140,160 | 2,140,160 | 2,140,160 | 2,140,160 |
| Final particle images (no.) | 135,694 | 84,814 | 238,580 | 48,169 |
| Map resolution (Å)<br>FSC threshold 0.143 | 2.6 | 2.6 | 2.2 | 2.9 |
| Helical rise (Å) | 4.85 | 4.84 | 4.82 | 4.82 |
| Helical twist (°) | -0.76 | -0.78 | -1.64 | -1.15 |
| **Refinement** |  |  |  |  |
| Model resolution (Å)<br>FSC threshold 0.5 | 3.1 | 2.8 | 2.7 | 3.2 |
| Map sharpening *B* factor (Å<sup>2</sup>) | -33 | -42 | -35 | -43 |
| Model composition |  |  |  |  |
| Non-hydrogen atoms | 2115 | 4175 | 1770 | 3408 |
| Protein residues | 279 | 552 | 219 | 438 |
| Ligands | 0 | 0 | 0 | 0 |
| *B* factors (Å<sup>2</sup>) |  |  |  |  |
| Protein | 59.1 | 58.8 | 96.5 | 96.47 |
| R.m.s. deviations |  |  |  |  |
| Bond lengths (Å) | 0.011 | 0.011 | 0.010 | 0.011 |
| Bond angles (°) | 2.154 | 2.008 | 1.927 | 1.965 |
| Validation |  |  |  |  |
| MolProbity score | 1.13 | 1.23 | 0.5 | 0.77 |
| Clashscore | 0.00 | 0.47 | 0.00 | 0.00 |
| Poor rotamers (%) | 0.00 | 0.21 | 0.00 | 0.00 |

## References

1. Goedert, M., Spillantini, M.G., Jakes, R., Rutherford, D., and Crowther, R.A. (1989). Multiple isoforms of human microtubule-associated protein tau: sequences and localization in neurofibrillary tangles of Alzheimer’s disease. Neuron 3, 519–526.

2. Scheres, S.H.W., Ryskeldi-Falcon, B., and Goedert, M. (2023). Molecular pathology of neurodegenerative diseases by cryo-EM of amyloids. Nature 621, 701–710.

3. Poorkaj, P., Bird, T.D., Wijsman, E., Nemens, E., Garruto, R.M., Anderson, L., Andreadis, A., Wiederholt, W.C., Raskind, M., and Schellenberg, G.D. (1998). Tau is a candidate gene for chromosome 17 frontotemporal dementia. Ann. Neurol. 43, 815–825.

4. Hutton, M., Lendon, C.L., Rizzu, P., Baker, M., Froelich, S., Houlden, H., Pickering-Brown, S., Chakraverty, S., Isaacs, A., Grover, A., et al. (1998). Association of missense and 5’-splicesite mutations in tau with the inherited dementia FTDP-17. Nature 393, 702–705.

5. Spillantini, M.G., Murrell, J.R., Goedert, M., Farlow, M.R., Klug, A., and Ghetti, B. (1998). Mutation in the tau gene in familial multiple system tauopathy with presenile dementia. Proc. Natl. Acad. Sci. USA 95, 7737–7741.

6. Kovacs, G.G., Ghetti, B., and Goedert, M. (2024) Tau proteinopathies. In: Greenfield’s Neuropathology, 10th Edition (C. Smith, A. Perry, G.G. Kovacs, T.S. Jacques eds), pp. 1073–1144. CRC Press, Boca Raton.

7. Schweighauser, M., Garringer, H.J., Klingstedt, T., Nilsson, K.P.R., Masuda-Suzukake, M., Murrell, J.R., Risacher, S.L., Vidal, R., Scheres, S.H.W., Goedert, M., et al. (2023). Mutation ΔK281 in MAPT causes Pick’s disease. Acta Neuropathol. 146, 211–226.

8. Shi, Y., Zhang, W., Yang, Y., Murzin, A.G., Falcon, B., Kotecha, A., van Beers, M., Tarutani, A., Kametani, F., Garringer, H.J., et al. (2021). Structure-based classification of tauopathies. Nature 598, 359–363.

9. Qi, C., Lövestam, S., Murzin, A.G., Peak-Chew, S., Franco, C., Bogdani, M., Latimer, C., Murrell, J.R., Cullinane, P.W., et al. (2025). Tau filaments with the Alzheimer fold in human MAPT mutants V337M and R406W. Nature Struct. Mol. Biol. 32, 1297–1304.

10. Schweighauser, M., Shi, Y., Murzin, A.G., Garringer, H.J., Vidal, R., Murrell, J.R., Erro, M.E., Seelaar, H., Ferrer, I., van Swieten, J.C., et al. (2025). Distinct tau filament folds in human MAPT mutants P301L and P301T. Nature Struct. Mol. Biol. 32, 1470–1478.

11. Pan, H.S., Merz, G.E., Li, A.N., Le, M.Q., Jo, H., Quddus, A., Yung, A., Kormos, R.C., Melo, A.A., Ramos, E.M., et al. (2026). Distinct tau filament folds in frontotemporal dementia due to the MAPT S305I mutation. BioRxiv 2026.02.12.705620.

12. Lövestam, S., Koh, F.A., van Knippenberg, B., Kotecha, A., Murzin, A.G., Goedert, M., and Scheres, S.H.W. (2022). Assembly of recombinant tau into filaments identical to those of Alzheimer’s disease and chronic traumatic encephalopathy. eLife 11, e76494.

13. Lövestam, S., Li, D., Wagstaff, J.L., Kotecha, A., Kimanius, D., McLaughlin, S.H., Murzin, A.G., Freund, S.M.V., Goedert, M., and Scheres, S.H.W. (2024) Disease-specific tau filaments assemble via polymorphic intermediates. Nature 625, 119–125.

14. Lövestam, S., Wagstaff, J.L., Katsinelos, T., Shi, J., Freund, S.M.V., Goedert, M., and Scheres, S.H.W. (2025) Twelve phosphomimetic mutations induce the assembly of recombinant fulllength human tau into paired helical filaments. eLife 14, RP104778.

15. Falcon, B., Zhang, W., Murzin, A.G., Murshudov, G., Garringer, H.J., Vidal, R., Crowther, R.A., Ghetti, B., Scheres, S.H.W., and Goedert, M. (2018). Structures of filaments from Pick’s disease reveal a novel tau protein fold. Nature 561, 137–140.

16. Shafei, R., Woollacott, I.O.C., Mummery, C.J., Bocchetta, M., Guerreiro, R., Bras, J., Warren, J.D., Lashley, T., Jaunmuktane, Z., and Rohrer, J.D. (2019). Two pathologically confirmed cases of novel mutations in the MAPT gene causing frontotemporal dementia. (2019). Neurobiol. Aging 87, 141.e15-141.e20.

17. Sanders, J., Schenk, V.W.D., and van Veen, P. (1939). A family with Pick’s disease. Verhandelingen der Koninklijke Nederlandse Akademie van Wetenschappen (tweede sectie) 38, 1–124.

18. Schenk, V.W. (1959). Re-examination of a family with Pick’s disease. Ann. Hum. Genet. 23, 325–333.

19. Groen, J.J., and Endtz, L.J. (1982). Hereditary Pick’s disease: second re-examination of the large family and discussion of other hereditary cases, with particular reference to electroencephalography, a computerized tomography. Brain 105, 443–459.

20. Spillantini, M.G., Crowther, R.A., Kamphorst, W., Heutink, P., and van Swieten, J.C. (1998). Tau pathology in two Dutch families with mutations in the microtubule-binding region of tau. Am. J. Pathol. 153, 1359–1363.

21. Bronner, I.F., ter Meulen, B.C., Azmani, A., Severijnen, L.A., Willemsen, R., Kamphorst, W., Ravid, R., Heutink, P., and van Swieten, J.C. (2005). Hereditary Pick’s disease with the G272V tau mutation shows predominant three-repeat tau pathology. Brain 128, 2645–2653.

22. Giannini, L.A.A., Ohm, D.T., Rozemuller A.J.M., Dratch, L., Suh, E., van Deerlin, V.M., Trojanowski, J.Q., Lee, E.B., van Swieten, J.C., Grossman, M., et al. (2022). Isoform-specific patterns of tau burden and neuronal degeneration in MAPT-associated frontotemporal lobar degeneration. Acta Neuropathol. 144, 1065–1084.

23. Probst, A., Tolnay, M., Langui, D., Goedert, M., and Spillantini, M.G. (1996). Pick’s disease: Hyperphosphorylated tau protein segregates to the somatoaxonal compartment. Acta Neuropathol. 92, 588–596.

24. Rosso, S.M., van Herpen, E., Deelen, W., Kamphorst, W., Severijnen, L.A., Willemsen, R., Ravid, R., Niermeijer, M.F., Dooijes, D., Smith, M.J., et al. (2002). A novel tau mutation, S320F, causes a tauopathy with inclusions similar to those in Pick’s disease. Ann. Neurol. 51, 373–376.

25. D’Souza, I., and Schellenberg, G.D. (2000). Determinants of 4-repeat tau expression. J. Biol. Chem. 275, 17700–17709.

26. Hasegawa, M., Smith, M.J., and Goedert, M. (1998). Tau proteins with FTDP-17 mutations have a reduced ability to promote microtubule assembly. FEBS Lett. 437, 207–210.

27. Chen, D., Bali, S., Singh, R., Wosztyl, A., Mullapudi, V., Vaquer-Alicea, J., Jayan, P., Melhem, S., Seelaar, H., van Swieten, J.C., et al. (2023). FTD-tau S320F mutation stabilizes local structure and allosterically promotes amyloid motif-dependent aggregation. Nature Commun. 14, 1625.

28. Von Bergen, M., Friedhoff, P., Biernat, J., Heberle, J., Mandelkow, E.M., and Mandelkow, E. (2000). Assembly of tau protein into Alzheimer paired helical filaments depends on a local sequence motif ((306)VQIVYK(311)) forming beta structure. Proc. Natl. Acad. Sci. USA 97, 5129–5134.

29. Macdonald, J.A., Bronner, I.F., Drynan, L., Fan, J., Curry, A., Fraser, G., Lavenir, I., and Goedert, M. (2019). Assembly of transgenic human P301S tau is necessary for neurodegeneration in murine spinal cord. Acta Neuropathol. Commun. 7, 44.

30. Goedert, M., Eisenberg, D.S., and Crowther, R.A. (2017). Propagation of tau aggregates and neurodegeneration. Annu. Rev. Neurosci. 40, 189–210.

31. Fitzpatrick, A.W.P., Falcon, B., He, S., Murzin, A.G., Murshudov, G., Garringer, H.J., Crowther, R.A., Ghetti, B., Goedert, M., and Scheres, S.H.W. (2017). Cryo-EM structures of tau filaments from Alzheimer’s disease brain. Nature 547, 185–190.

32. Tarutani, A., Arai, T., Murayami, S., Hisanaga, S.I., and Hasegawa, M. (2018). Potent prion-like behaviors of pathogenic alpha-synuclein and evaluation of inactivation methods. Acta Neuropathol. Commun. 6, 29.

33. Garcia-Nafria, J., Watson, J., and Greger, I. (2016). IVA cloning: A single-tube universal cloning system exploiting bacterial in vivo assembly. Sci. Rep. 6, 27459.

34. Scheres, S.H.W. (2012). A Bayesian view on cryo-EM structure determination. J. Mol. Biol. 415, 406–418.

35. Kimanius, D., Dong, L., Sharov, G., Nakane, T., and Scheres, S.H.W. (2021). New tools for automated cryo-EM single-particle analysis in RELION-4.0. Biochem. J. 478, 4169–4185.

36. He, S., and Scheres, S.H.W. (2017). Helical reconstruction in RELION. J. Struct. Biol. 198, 163–176.

37. Zivanov, J., Nakane, T., Forsberg, B.O., Kimanius, D., Hagen, W.J., Lindahl, E., and Scheres, S.H.W. (2018). New tools for autyomated high-resolution cryo-EM structure determination in RELION-3. eLife 7, e42166.

38. Rohou, A., and Grigorieff, N. (2015). CTFFIND4: Fast and accurate defocus estimation from electron micrographs. J. Struct. Biol. 192, 216–221.

39. Kimanius, D., Jamali, K., Wilkinson, M.E., Lövestam, S., Velazhahan, V., Nakane, T., and Scheres, S.H.W. (2024). Data-driven regularization lowers the size barrier of cryo-EM structure determination. Nature Meth. 21, 1216–1221.

40. Zivanov, J., Nakane, T., Scheres, S.H.W., 2020. Estimation of high-order aberrations and anisotropic magnification from cryo-EM data sets in RELION-3.1. IUCrJ 7, 253–267.

41. Zivanov, J., Nakane, T., Scheres, S.H.W., 2019. A Bayesian approach to beam-induced motion correction in cryo-EM single-particle analysis. IUCrJ 6, 5–17.

42. Scheres, S.H.W., and Chen, S. (2012). Prevention of overfitting in cryo-EM structure determination. Nature Meth. 9, 853–854.

43. Emsley, P., Lohkamp, B., Scott, W.G., and Cowtan, K. (2010). Features and development of Coot. Acta Crystallogr. D 66, 486–501.

44. Falcon, B., Zivanov, J., Zhang, W., Murzin, A.G., Garringer, H.J., Vidal, R., Crowther, R.A., Newell, K.L., Ghetti, B., Goedert, M., et al. (2019). Novel tau filament fold in chronic traumatic encephalopathy encloses hydrophobic molecules. Nature 568, 420–423.

45. Yamashita, K., Palmer, C.M., Burnley, T., and Murshudov, G.N. (2021). Cryo-EM single-particle structure refinement and map calculation using SERVALCAT. Acta Crystallogr. D 77, 1282–1291.

46. Croll, T.I. (2018). ISOLDE: A physically realistic environment for model building into low-resolution electron-density maps. Acta Crystallogr. D 74, 519–530.

47. Murshudov, G.N., Vagin, A.A., and Dodson, E.J. (1997). Refinement of macromolecular structures by the maximum-likelihood method. Acta Crystallogr. D 53, 240–255.

48. Murshudov, G.N., Skubák, P., Lebedev, A., Pannu, N.S., Steiner, R.A., Nicholls, R.A., Winn, M.D., Long, F., and Vagin, A.A. (2011). REFMAC5 for the refinement of macromolecular crystal structures. Acta Crystallogr. D 67, 355–367.

49. Chen, V.B., Arendal III, W.B., Headd, J.J., Keedy, D.A., Immormino, R.M., Kapral, G.J., Murray, L.W., Richardson, J.S., and Richardson, D.C. (2010). MolProbity: All-atom structure validation for macromolecular crystallography. Acta Crystallogr. D 66, 12–21.

50. Pettersen, E.F., Goddard T.D., Huang, C.C., Meng, E.C., Couch, G.S., Croll, T.I., Morris, J.H., and Ferrin, T.E. (2021). UCSF ChimeraX: Structure visualization for researchers, educators and developers. Prot. Sci. 30, 70–82.

51. Schrödinger, L.L.C. (2015). The PyMOL Molecular Graphics System, Version 1.8. Schrödinger, LLC.

52. Scheres, S.H.W. (2026) The amyloid packing difference: a pairwise comparison metric for amyloid structures. BioRxiv 2026.02.18.706523.

